# The DEAD-box RNA helicase Ded1 from yeast is associated with the signal recognition particle (SRP), and its enzymatic activity is regulated by SRP21

**DOI:** 10.1101/2020.11.08.373522

**Authors:** Hilal Yeter-Alat, Naïma Belgareh-Touzé, Emmeline Huvelle, Molka Mokdadi, Josette Banroques, N. Kyle Tanner

## Abstract

The DEAD-box RNA helicase Ded1 is an essential yeast protein involved in translation initiation. It belongs to the DDX3 subfamily of proteins implicated in developmental and cell-cycle regulation. *In vitro*, the purified Ded1 protein is an ATP-dependent RNA binding protein and an RNA-dependent ATPase, but it lacks RNA substrate specificity and enzymatic regulation. Here we demonstrate by yeast genetics, *in situ* localization and *in vitro* biochemical approaches that Ded1 is associated with, and regulated by, the signal recognition particle (SRP), which is a universally conserved ribonucleoprotein complex required for the co-translational translocation of polypeptides into the endoplasmic reticulum lumen and membrane. Ded1 is physically associated with SRP components *in vivo* and *in vitro*. Ded1 is genetically linked with SRP proteins. Finally, the enzymatic activity of Ded1 is inhibited by SRP21 with SCR1 RNA. We propose a model where Ded1 actively participates in the translocation of proteins during translation. Our results open a new comprehension of the cellular role of Ded1 during translation.

## INTRODUCTION

The DEAD-box family of RNA helicases are ubiquitous proteins found in all three kingdoms of life, and they are implicated in all processes involving RNA, from transcription, splicing, ribosomal biogenesis, RNA export, translation to RNA decay [reviewed by (1–3)]. They belong to the DExD/H superfamily 2 (SF2) of putative RNA and DNA helicases that contain catalytic cores consisting of two, linked, RecA-like domains containing conserved motifs associated with ligand binding and NTPase activity, where the majority of the proteins are ATPases. In addition, they often contain highly variable amino- and carboxyl-terminal domains [reviewed by (4,5)]. The DEAD-box proteins are ATP-dependent RNA binding proteins and RNA-dependent ATPases that have been shown to remodel RNA and ribonucleoprotein (RNP) complexes and to unwind short RNA duplexes *in vitro*, but they are not processive, and they generally have shown little or no substrate specificity (1–3). However, recent single-molecule studies of the DEAD-box protein Ded1 indicate that these properties may be secondary to their ability to form ATP-dependent clamps on RNA (6).

A number of crystal structures of DEAD-box proteins have been solved in the presence and absence of ligands [reviewed by (7)]. In the absence of ATP, the two RecA-like domains are unconstrained (“open” conformation) and the proteins have low affinity for RNA. In the presence of ATP, the two RecA-like domains are highly constrained (“closed” conformation) and have a high affinity for the RNA. RecA-like domain 2 binds the 5’ end of the RNA as a single strand in the form of an A helix. In contrast, RecA-like domain 1 binds the 3’ end of the RNA with a kink as a result of steric hindrance from residues from motifs Ib and GG that is incompatible with a duplex. This is considered the mechanism for the duplex unwinding activity, but it also effectively locks the protein onto the RNA and prevents sliding. Indeed, Ded1 in the presence of the nonhydrolyzable ATP analog ADP-BeF_x_ forms long-lived complexes on RNA *in vitro* (8).

Ded1 is a budding-yeast DEAD-box protein that is the functional homolog of mammalian DDX3 [reviewed by (9–12)]. It is an essential gene in *Saccharomyces cerevisiae* that can be rescued by the expression of its orthologs from other eukaryotes, including human DDX3 [(13) and references therein]. Thus, the functional activity of Ded1 is conserved throughout eukaryotes. Ded1 is considered a general translation-initiation factor that is important for 43S ribosome scanning to the initiation codon and formation of the 48S complex at the AUG codon [(14–16) and references therein]. We have shown that Ded1 is a cap-associated factor that actively shuttles between the nucleus and cytoplasm using both the XpoI/Crm1 and Mex67/TAP nuclear pore complexes (13). Moreover, it interacts with both the nuclear and cytoplasmic 3’ polyA-binding proteins Nab2 and Pab1, respectively. We found that theses cap-associated factors stimulate the RNA-dependent ATPase activity of Ded1 (13). The activity of Ded1 is also modulated by Gle1 and by the Xpo1-Ran[GTP] complex (13,17,18). Other work has shown that Ded1 is sequestered in cytoplasmic foci (P-bodies or stress granules) with translation inactive mRNAs during conditions of stress [reviewed by (19–22)]. Human DDX3 has similar properties [reviewed by (23)].

We are interested in better understanding the role of Ded1 in the cell. To this end, we used a modified photoactivable-ribonucleoside-enhanced-crosslinking-and-immunoprecipitation (PAR-CLIP) technique to identify RNA substrates of Ded1 *in vivo* (24,25). We identified the Small Cytoplasmic RNA 1 (SCR1) as a major noncoding RNA that crosslinked to Ded1. SCR1 is the RNA component of the signal recognition particle (SRP) that is important for the co-translational translocation of polypeptides into the lumen and membrane of the endoplasmic reticulum [ER; reviewed by (26–29)]. SRP-dependent translation is conserved across all organisms, from prokaryotes to eukaryotes, including a highly reduced version for chloroplasts (30–32). It seemed possible that Ded1 was implicated in SRP-dependent translation.

SRP-dependent translation is a complicated and multi-step process that is still incompletely understood and that involves a number of still controversial elements. In eukaryotes, the SRP consists of the noncoding RNA (7S or 7SL in metazoans) and six SRP proteins (SRP9, SRP14, SRP19, SRP54, SRP68 and SRP72). The SRP RNA consists of two functional elements called the Alu and S domains (33,34). The yeast SRP complex consists of the SCR1 RNA and the equivalent proteins except Sec65 substitutes for the smaller SRP19 and the novel SRP21 protein replaces SRP9 [reviewed by (35)]. Moreover, in yeast, SRP14 forms a homodimer on the Alu domain of SCR1, which is in contrast to the heterodimer of SRP14-SRP9 in metazoans (35–37). The role of the yeast SRP21 protein is unclear, although it is considered the structural homolog of SRP9 (38). SRP54 and Sec65 interact with the extremity of the S domain of SCR1, and SRP68 and SRP72 interact at the junction between the Alu and S domains (27,35). Yeast lacks the “classical” structure of the Alu domain that includes helices 3 and 4, and it contains additional hairpins between the Alu and S domains; it is about 75% bigger (33,34). The structure and role of these additional hairpins are largely unknown. Eukaryotes lack helix 1 that is found in prokaryotes.

In the classical interpretation, the SRP associates with the ribosome during translation when the SRP54 GTPase binds the hydrophobic signal peptide as it emerges from the exit channel of the 60S ribosomes as a ribosome nascent-chain complex [RNC; reviewed by (26–28)]. This causes the ribosomes to pause translation and permits the SRP-ribosome complex to associate with the SRP receptor (SR) on the ER that consists of the membrane associated SRP101 and the integral membrane protein SRP102 (SRα and SRβ, respectively, in metazoans). In yeast and metazoans, SRP14 and the Alu domain of the SRP RNA play an important role in this “pausing” by blocking the GTP-dependent elongation factor eEF2 from binding at the GTPase-associated center near the mRNA entry channel at the interface between the 40S and 60S ribosomes (37,39,40). The interactions with the ribosomes depend on the SRP proteins; human 7SL RNA does not bind the ribosomes by itself (41). The SRP-ribosome complex eventually associates with the Sec61 translocon on the ER membrane, the SRP dissociates from the ribosome and translation continues with the polypeptide inserted into the ER lumen or membrane. Ded1 could be intimately associated with this process.

We find that Ded1 is an SRP-associated factor. Ded1 is genetically linked to the SRP proteins, and it associates with SRP complexes in pull-down experiments and sucrose gradients. The purified recombinant SRP proteins physically interact with Ded1 *in vitro*. Moreover, fluorescence microscopy shows that Ded1 is associated with mRNAs at the ER membrane in the cell. RNA binding assays show that Ded1 has a high affinity for SCR1 RNA. Finally, the ATPase activity of Ded1 is inhibited by SRP21, and it is inhibited much more when SRP21 is associated with SCR1 than with other RNAs. We propose a model where Ded1 plays an important role in the SRP-dependent translation of proteins.

## RESULTS

### Ded1 associated with SRP factors *in vivo*

We have been using a modified photoactivable-ribonucleoside-enhanced-crosslinking-and-immunoprecipitation (PAR-CLIP) technique to identify RNA substrates of Ded1 *in vivo* (24,25). This is an ongoing project, but in the process we recovered significant crosslinks to the noncoding RNA SCR1 that is part of the SRP involved in co-translational transport of polypeptides into the membrane and lumen of the ER (Figure 1). Although unanticipated, this result was consistent with our previous observations that Ded1 cosediments and associates with complexes containing SRP proteins on polysome sucrose gradients, as determined by pull-down experiments and mass spectroscopy analyses (13). These data show that Ded1 and SRP proteins SRP14, SRP21, SRP54 and SRP68 sediment at a position corresponding to ∼26S (Table 1). Moreover, SRP14, SRP21, Sec65 and SRP68 are in stable complexes associated with Ded1 on the sucrose gradients that are pulled down with Ded1-specific IgG (Table 2). These results indicated that Ded1 might be associated with ribosomes translating mRNAs encoding ER proteins.

**Figure 1.**
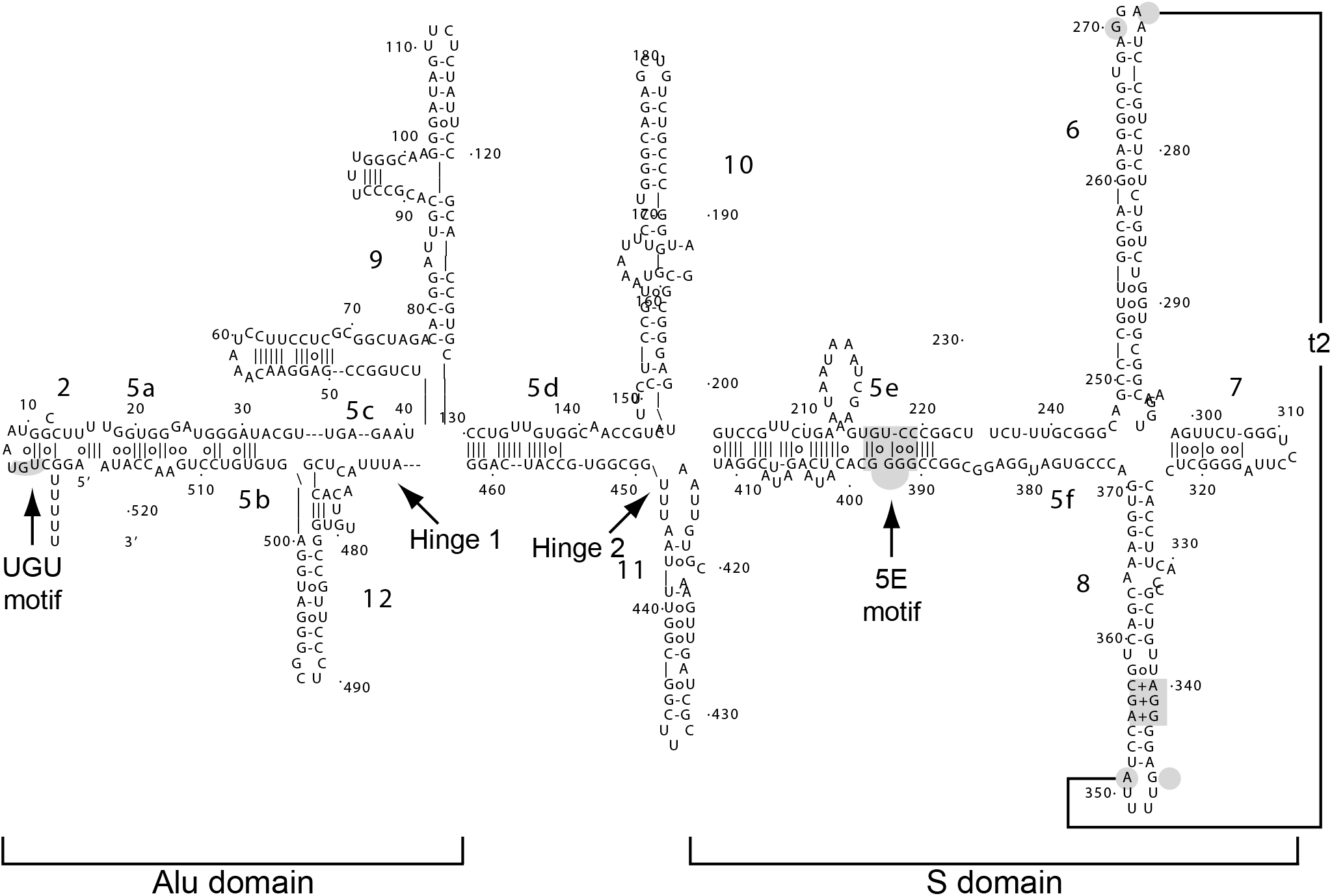
Secondary structure model of yeast SCR1 of Zwieb *et al.* The model is based on phylogenetically conserved features found in SRP RNAs and on structural probing experiments (33,34). Yeast and other fungal SRP RNAs are unusual in that they are much larger than in other organisms, and they lack the characteristic structure consisting of hairpins 3 and 4 of the Alu domain. Yeast has the additional hairpins 9, 10, 11 and 12 that are poorly characterized and that have other proposed secondary structures. Conserved sequence motifs and tertiary interactions are shown in gray.

**Table 1.** Sucrose gradients fractions; nano-LC ESI MS/MS analysis

| Protein<sup>b</sup> | Fraction(s)<sup>c</sup> | Meta Score<sup>d</sup> | #Spectres<sup>e</sup> | SC%<sup>f</sup> | RMS (ppm)<sup>g</sup> |
| --- | --- | --- | --- | --- | --- |
| Ded1 | 6 | 213.4 | 7 | 14.2 | 6.60 |
| Ded1 | 7 | 1455.7 | 50 | 50.8 | 8.04 |
| SRP14 | 7 | 80.7 | 4 | 13.0 | 5.25 |
| SRP21 | 7 | 255.5 | 10 | 28.7 | 7.30 |
| SRP54 | 7 | 229.4 | 6 | 8.9 | 6.40 |
| SRP68 | 7 | 88.2 | 2 | 2.2 | 8.01 |
| ENO2 | 6 | 5771.5 | 639 | 96.6 | 5.68 |
| ENO2 | 7 | 4741.5 | 359 | 94.7 | 6.50 |
| SSA2 | 6 | 4335.6 | 280 | 84.4 | 10.24 |
| SSA2 | 7 | 3999.2 | 226 | 74.5 | 8.37 |
| FBA1 | 6 | 2125.7 | 186 | 80.5 | 6.52 |
| FBA1 | 7 | 1653.2 | 116 | 79.7 | 6.32 |

<sup>a</sup> Data was collected and analyzed as previously described (13). In brief, 0.5 ml fractions were collected starting from the top of the gradient and subjected to nano-liquid-chromatography electron-spray mass spectrometry analysis <sup>b</sup> ENO2, SSA2 and FBA1 are reference proteins; they showed the strongest signals in the fraction at the top of gradient. <sup>c</sup> Fractions correspond to those shown in Supplemental Figure S7 of Senissar *et al.* (13). <sup>d</sup> Mascot probably-based scoring. <sup>e</sup> Spectral counting; the same peptide is fragmented up to six times over a mean elution time of 30 seconds. <sup>f</sup> Percentage of the protein sequence covered. <sup>g</sup> Mean error in ppm.

**Table 2.** Ded1-IgG Pull-down of sucrose gradients fractions; Nano-LC ESI MS/MS analysis

| Protein<sup>b</sup> | Fraction(s)<sup>c</sup> | Meta Score<sup>d</sup> | #Peptides<sup>e</sup> | SC%<sup>f</sup> | RMS (ppm)<sup>g</sup> |
| --- | --- | --- | --- | --- | --- |
| Ded1 | 6 | 413.4 | 25 | 40.6 | 3.51 |
| Ded1 | 7 | 368.2 | 8 | 15.6 | 1.73 |
| SRP14 | 6 | 387.0 | 9 | 51.4 | 2.54 |
| SRP14 | 7 | 88.2 | 2 | 13.0 | 2.32 |
| SRP21 | 6 | 385.5 | 6 | 50.3 | 3.15 |
| SRP21 | 7 | 95.4 | 2 | 13.2 | 1.99 |
| Sec65 | 6 | 345.1 | 7 | 31.9 | 2.69 |
| SRP68 | 6 | 66.4 | 1 | 2.2 | 0.94 |
| ENO2 | 6, 7 | — | — | — | — |
| SSA2 | 6 | 395.9 | 9 | 21.3 | 2.83 |
| SSA2 | 7 | — | — | — | — |
| FBA1 | 6, 7 | — | — | — | — |

<sup>a</sup> Data was collected and analyzed as previously described (13). Equivalent fractions as shown in Table 1 were subjected to IgG-Ded1 pull-downs with protein-A Sepharose beads and subjected to mass spectrometry analysis. <sup>b</sup> ENO2, SSA2 and FBA1 are reference proteins; they showed the strongest signals in Table 1. <sup>c</sup> Fractions correspond to those shown in Supplemental Figure S7 of Senissar *et al.* (13). <sup>d</sup> Mascot probably-based scoring. <sup>e</sup> Number of peptide fragments recovered. <sup>f</sup> Percentage of the protein sequence covered. <sup>g</sup> Mean error in ppm.

To elaborate on these observations, we did pull-down experiments of yeast extracts with IgG against Ded1 and then subjected the recovered material to Northern blot analysis with a ^32^P-labeled DNA probe against SCR1. We used a probe against PGK1 mRNA as a positive control because it also was found to crosslink efficiently to Ded1, and a probe against RPL20B mRNA as a negative control as we obtained little crosslinking on this RNA, even though it is a highly expressed mRNA (42). However, the resulting signals were insufficiently sensitive. This was not unexpected as Ded1 interacts with a large number of mRNAs, and the RNAs of interest represented a small fraction of these total RNAs (14,43). Hence, we performed RT-PCR on the samples using oligonucleotides specific for the three RNA. Both SCR1 and PGK1 RNAs were amplified much more in the fractions pulled down with Ded1-specific IgG than in the control fractions that were pulled down with pre-immune IgG (Figure 2A). In contrast, RPL20B mRNA was weakly amplified in both cases. Hence, Ded1 associated with SCR1 RNA *in vivo*.

**Figure 2.**
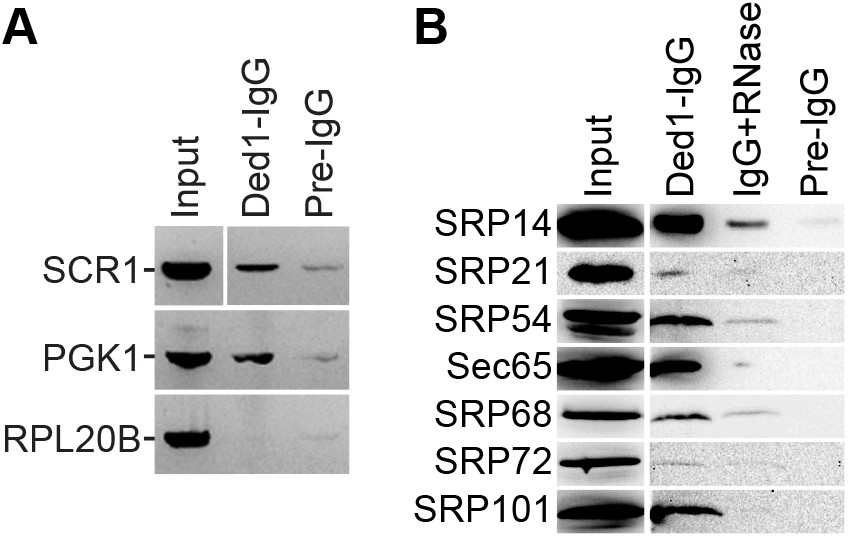
Ded1-IgG pull-downs of yeast extracts. Ded1-specific IgG (Ded1-IgG) or IgG from pre-immune serum (Pre-IgG) were used to recover the associated factors. Input, a fraction of the yeast extract used in the pull-down experiments was directly loaded onto the gel or RT-PCR amplified. (**A**) Purified RNA from yeast extracts (∼20% of input) or from IgG pull-downs was reverse transcribed and PCR amplified for 25 cycles with gene-specific oligonucleotides. The resulting products were electrophoretically separated on a 2% agarose gel containing ethidium bromide, and the products visualized with a Gel Doc XR+ (Bio-RAD). (**B**) Western blot analysis of HA-tagged SRP proteins. Proteins were electrophoretically separated on a 12% SDS-PAGE, transferred to nitrocellulose membranes, and then revealed with anti-HA IgG. Input, 40 µg (∼10%) of the yeast extract was directly loaded on the gel. Ded1-IgG, Ded1-specific IgG was used to pull-down Ded1 associated proteins. IgG+RNase, complexes bound to Ded1-IgG-protein A beads were digested with RNase A (1mg/ml) prior to washing and elution.

It was possible that Ded1 interacted with the SCR1 RNA independently of the SRP proteins. To test this, we did Ded1-IgG pull-down experiments, SDS-PAGE separation and Western blot analyses of the recovered proteins. However, we were only able to obtain antibodies against Sec65 [generously provided by Martin R. Pool; (44)]. (We later made IgG against SRP21.) Hence, we cloned all the *SRP* genes with amino-terminal HA tags, and we did Ded1-IgG pull-downs with strains independently expressing each tagged protein (Figure 2B). We recovered a significant amount of SRP14, SRP54, Sec65 and SRP101. Likewise, we digested the Ded1-IgG-bound complexes with RNase A prior to elution to determine if these complexes depended on SCR1 RNA; in all cases, the signals for the SRP proteins were reduced, but the signals were still more than for the pre-immune-IgG control (Figure 2B). Oddly, we detected little HA-tagged SRP21 even though it was prominent in our previous mass spectrometry analyses [Tables 1 and 2; (13)]. Moreover, we detected little HA-SRP21 in yeast extracts even though it is of similar size and has similar expression levels as SRP14 (Figure 2B; Table 3). It was possible that the HA tag increased the proteolytic degradation of the protein during extraction or that the HA tag by itself was proteolytically removed. Thus, these results showed that Ded1 interacted with complexes containing the SRP proteins, and that these complexes were stabilized by SCR1 RNA (Figure 2B). Notably, the bound complexes also contained the SR receptor SRP101, which binds with SRP complexes associated with the ER during translation (26,35). We did not detect SRP102, but it contains an integral membrane domain that could limit its recovery. Thus, Ded1 physically associated with the SRP complex *in vivo*.

**Table 3.** Protein characteristics

| Protein | Gene | Length (aa) | MW (gm/mole) | pK<sub>i</sub> | Abundance<sup>b</sup> | Half life (hr)<sup>c</sup> |
| --- | --- | --- | --- | --- | --- | --- |
| Ded1 | YOR204W | 604 | 65554.7 | 7.98 | 25034 ± 5043 | 9.1 |
| SRP14 | YDL092W | 146 | 16442.1 | 10.38 | 5858 ± 1306 | 9.6 |
| SRP21 | YKL122C | 167 | 18451.7 | 11.14 | 4248 ± 1211 | 7.9 |
| SRP54 | YPR088C | 541 | 59630.3 | 9.04 | 8411 ± 2308 | 7.0 |
| Sec65 | YML105C | 273 | 31177.3 | 9.62 | 5632 ± 2232 | 9.7 |
| SRP68 | YPL243W | 599 | 69014.7 | 9.15 | 8214 ± 1962 | 9.9 |
| SRP72 | YPL210C | 640 | 73568.8 | 10.0 | 6363 ± 1828 | 9.4 |
| SRP101 | YDR292C | 621 | 69276.8 | 7.19 | 4444 ± 2444 | 9.9 |
| SRP102 | YKL154W | 244 | 26975.3 | 8.02 | 5274 ± 1935 | 10.1 |

<sup>a</sup> Data taken from https://www.yeastgenome.org.
<sup>b</sup> Median abundance (molecules/cell).
<sup>c</sup> Data from (97).

### Ded1 cosedimented with SRP factors

In our previous work, we found that Ded1 migrated at a position corresponding to ∼26S on sucrose gradients, but the Ded1-containing fractions were only partially resolved from the protein peak at the top of the gradient (13). Hence, we modified the previous sucrose gradient conditions to better resolve the different complexes. In both cases, we used 5 mM MgCl_2_ because Ded1 was found to dissociate from higher molecular weight complexes at higher Mg^2+^ concentrations and sediment within the protein peak at the very top of gradient. We did sucrose gradients of yeast strains independently expressing each HA-tagged SRP protein under identical conditions.

The polysome profile (Figure 3A) showed well resolved ribosome peaks that corresponded to the expected distribution of ribosomal RNAs (Figure 3B). Northern blot analyses with a SCR1-specific ^32^P-labeled probe showed that the vast majority of the SCR1 RNA migrated as a narrow peak centered at fraction 4 (arrow), but RNA was detected throughout the gradient (Figure 3C). Western blot analyses showed a very heterogeneous distribution of the proteins (Figure 3D). Ded1 and Sec65 were concentrated near the top of the gradients while the other SRP proteins were more widely distributed. SRP54 migrated as two bands that probably represented different modified forms of the protein. Oddly, we did not detect HA-tagged SRP21 even though it was prominent in our previous mass spectrometry analyses [Tables 1 & 2; (13)]; however, this result was consistent with the IgG pull-down experiments. Moreover, all of the detected proteins were found in fraction 4 that had the most SCR1 RNA.

**Figure 3.**
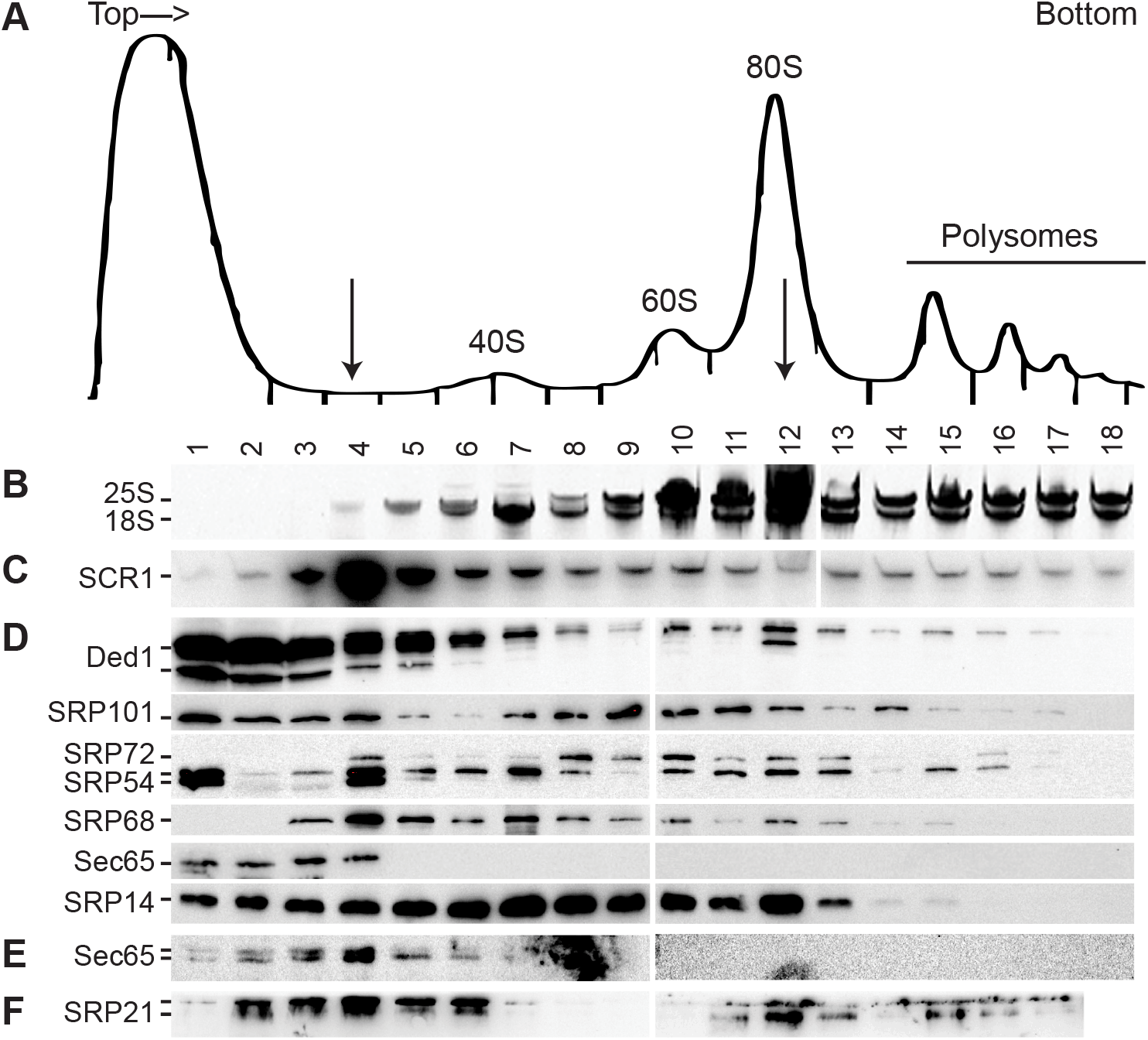
Polysome sucrose gradient of cell extracts. Extracts of cell cultures individually expressing HA-tagged proteins were treated with cycloheximide and separated on 10–50% sucrose gradients. Note that conditions were modified from those previously used to better separate lower molecular-weight complexes (13). (**A**) Trace of a representative sucrose gradient monitored spectroscopically at 254 nm with 0.5 ml fractions collected from the top of the gradient. (**B**) Extracted RNAs from different fractions were electrophoretically separated on a 6% polyacrylamide gel containing 7 M urea and the separated RNAs were visualized with ethidium bromide. (**C**) Northern blot analysis of the material separated as shown in B and transferred to nitrocellulose membranes. The RNA was detected with a ^32^P-labeled DNA oligonucleotide specific for SCR1. (**D**) Western blot analysis of HA-tagged proteins. Precipitated proteins from different fractions were electrophoretically separated on a 12% SDS-PAGE, transferred to nitrocellulose membranes and probed with IgG against the HA tag and Ded1. (**E**) Western blot analysis of endogenous Sec65 probed with IgG against Sec65. (**F**) Western blot analysis of endogenous SRP21 probed with IgG against SRP21.

It is known that constitutive over-expression of the SRP proteins can cause them to accumulate in the nucleus, which may affect their distribution on the sucrose gradients (45). Consequently, we did sucrose gradients of the endogenously-expressed Sec65 and probed the membranes with Sec65-specific IgG; it showed a similar distribution as the HA-tagged protein, but it was present as a doublet (Figure 3E). Likewise, we made SRP21-specific IgG to detect endogenous SRP21 in the gradients (Figure 3F). The vast majority of the protein was stuck in the well of the gel or migrated only a short distance into the gel that indicated that SRP21 formed large, partially insoluble aggregates. We obtained similar results with recombinant SRP21 in the presence of RNA (see below section: **SRP21 did not block Ded1 binding to SCR1**). Nevertheless, the majority of both Sec65 and SRP21 sedimented at the position corresponding to the bulk of the SCR1 RNA in the gradients.

The data were consistent with Ded1 interacting with the SRP complex. However, the vast majority of the material was not associated with translating ribosomes, although there was a smaller peak of Ded1 and most of the SRP proteins in fraction 12 that corresponded to the 80S complex (arrow). These results indicated either that most of Ded1 and the SRP complex were not actively involved in translation or that the complexes were not stably associated with the ribosomes under the conditions used. Thus, it was unclear as to the functional role of the Ded1-SRP interactions.

### Ded1 was genetically linked to SRP proteins

We next tested to see if there was a genetic link between Ded1 and the SRP proteins as we previously demonstrated for the nuclear and cytoplasmic cap-associated proteins (13). We used the same cold-sensitive mutant F162C of Ded1 in the *ded1::HIS3* deletion strain and over-expressed the SRP proteins and SCR1 RNA from the pMW292 and pM299 plasmids (44,46). Liquid cultures were serially diluted and spotted on 5-FOA plates that were incubated at 18°C, 30°C and 36°C (Figure 4). The results showed a slight enhancement of growth at 18°C that was consistent with a genetic interaction between Ded1 and the SRP complex, but the signal was too weak to demonstrate a clear link. The weak multicopy suppression was not unexpected because Ded1 is implicated in the expression of multiple mRNAs that are not associated with SRP complexes (14,43); expression of these mRNAs would be insensitive to the over-expressed SRP factors. Thus, it was possible that the SRP complex would be more sensitive to the level of Ded1 expression than vice versa.

**Figure 4.**
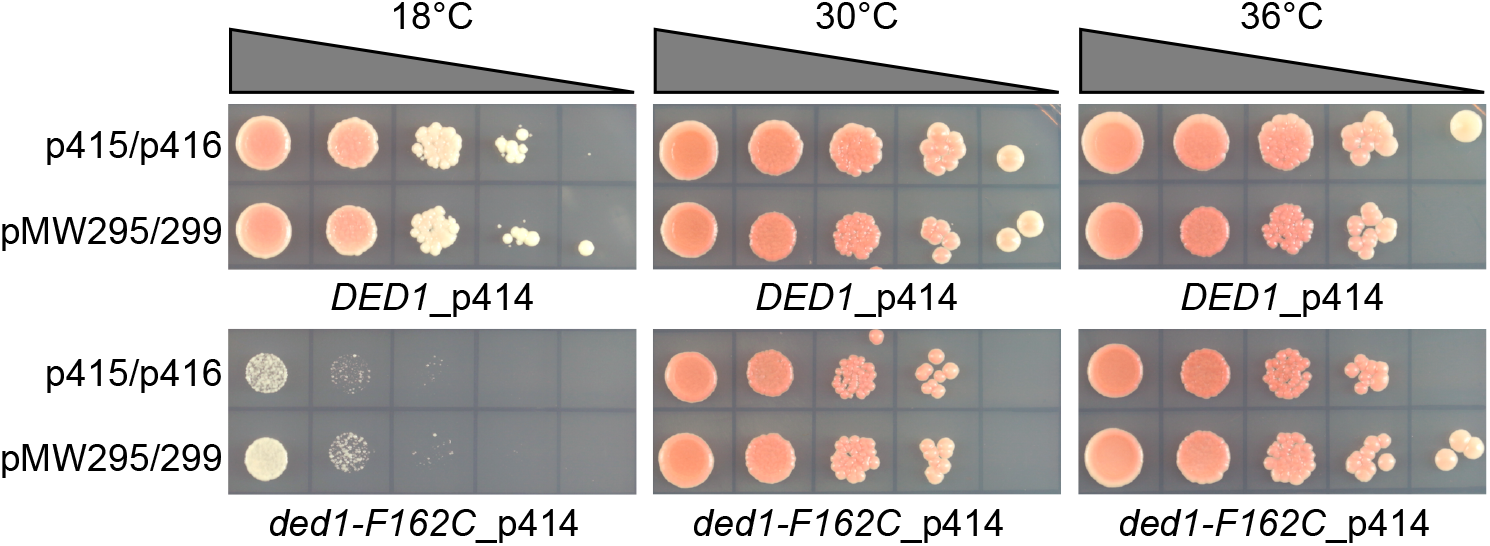
Multiple-copy suppression of the *ded1-F162C* cold-sensitive phenotype. The *ded1::HIS3* deletion strain containing the *DED1 URA* plasmid were transformed with plasmids expressing Ded1 wildtype or Ded1-F162C mutant proteins. They were subsequently transformed with pMW295 and pMW299 plasmids expressing the SRP components or with the empty plasmids. Liquid cultures were then serial diluted by a factor of 10 and spotted on synthetic defined (SD) medium plates containing 5-FOA and incubated for 3 days at 30°C and 36°C and for 5 days at 18°C

Previous work has shown that loss of any SRP component leads to a slow-growth phenotype (47–49), although yeast cells are eventually able to adapt to this loss (50,51). Thus, we obtained yeast strains with the *DED1*, *SRP14*, *SRP21*, *SEC65*, *SRP68, SRP72* and *SRP101* genes under the control of a tetracycline-regulated promoter that could be suppressed with doxycycline (52). Unfortunately, SRP54 under the tetracycline promoter was not available. Cultures of the different strains were grown in liquid culture, serially diluted and spotted on agar plates in the presence or absence of 10 µg/ml of doxycycline. All the strains except SRP101 showed reduced growth with the constitutive expression of the proteins, which was most apparent at 18°C (Figure 5). In the presence of doxycycline, all the strains show strongly reduced growth except SRP21, Sec65 and SRP101, which showed a slight reduction. Western blot analysis of liquid cultures of *TET-SRP21* and *TET-SEC65* grown for up to 24 h in the presence of 10 µg/ml of doxycycline showed no diminution in protein level when probed with SRP21-IgG or Sec65-IgG, respectively (data not shown). This indicated that either the *TET* promoter was not completely shut down with doxycycline in these strains or that the proteins were particularly stable. Interestingly, *TET-SRP21* actually grew slightly better in the presence of doxycyline, which further indicated that constitutive expression of the proteins was detrimental (Figure 5). Constitutive and overexpression of Ded1 was previously shown to inhibit cell growth (53,54).

**Figure 5.**
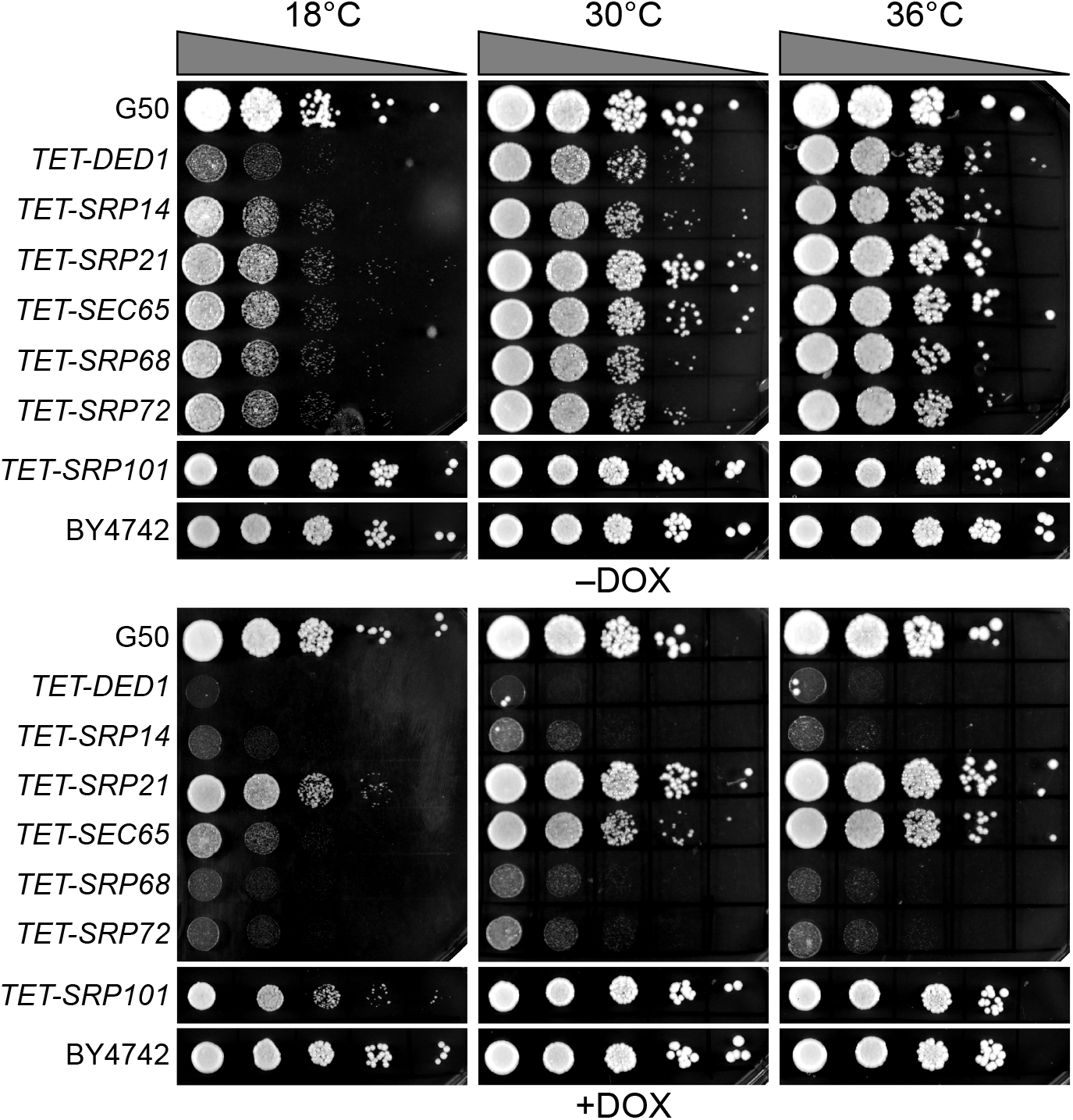
Phenotypes of proteins expressed with tetracycline promoters. Liquid cultures of the indicated strains were serially diluted by a factor of 10 and plated on YPD (yeast extract, peptone, dextrose) rich-medium agar plates, except for *TET-SRP101* and BY4742 that were plated on SD medium agar plates, in the presence (+DOX) or absence (-DOX) of 10 µg/ml of doxycycline. The G50 and BY4742 strains show wildtype growth. Plates were incubated for 2 days at 30°C and 36°C, and for 4 days at 18°C for the YPD plates, and for 4 days and 7 days, respectively, for the SD plates.

We transformed the different *TET* strains with a plasmid containing *DED1* under control of the very strong *GPD* promoter and compared it with cells transformed with the empty plasmid and with wildtype yeast cells (55). We likewise transformed the cells with a plasmid expressing the mutant Ded1-F162C protein that had reduced ATP binding and enzymatic activity (46). As expected, the yeast stains *TET-SRP21* and *TET-SEC65*, which continued to express the SRP proteins, showed reduced growth on the plates due to the inhibitory effects of the overexpressed Ded1 (Figure 6). In contrast, SRP14, SRP68 and SRP72 showed enhanced growth despite the inhibitory effects of Ded1 (Figure 6). The Ded1-F162C mutant showed little or no stimulatory effect, which indicated that the enzymatic function of Ded1 was important for the enhanced growth. Thus, high expression of Ded1 partially suppressed the slow-growth phenotype of strains depleted for SRP proteins, and this result established a genetic link between Ded1 and the SRP complex.

**Figure 6.**
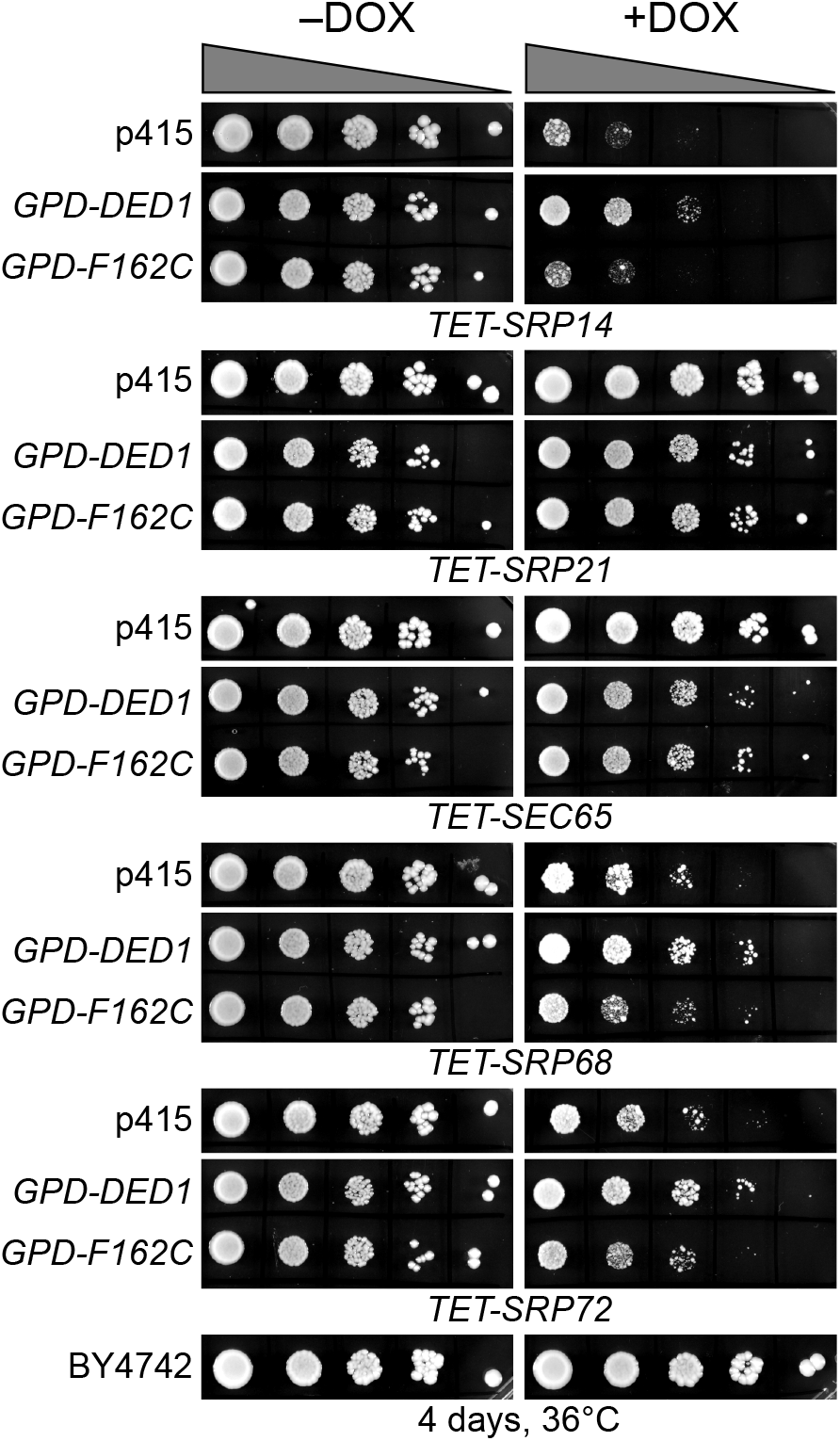
Ded1 multicopy suppression of SRP protein depletions. Cells of the indicated strains with the *TET* promoter were grown in SD-LEU medium, serially diluted by a factor of 10 and spotted on SD-LEU agar plates with (+DOX) or without (-DOX) 10 µg/ml of doxycycline. Cultures were grown 4 days at 36°C. p415, empty *LEU* plasmid; *GPD-DED1*, Ded1 in p415 with the high expression *GPD* promoter and *CYC1* terminator; *GPD-F162C*, a Ded1 mutant with reduced ATP binding and enzymatic activity (46). BY4742, a wildtype yeast strain showing unimpeded growth. The phenotypes were most apparent at 36°C, but similar effects were obtained at 30°C.

### Ded1 was in cellular foci associated with the endoplasmic reticulum

The next question we asked was whether Ded1 co-localized with the ER as would be expected if it was associated with SRP-ribosome complexes that were translating mRNAs encoding polypeptides translocated into the ER. However, we and others have shown that Ded1 has a diffuse location within the cytoplasm under normal growth conditions (13,53,56). Nevertheless, some of the Ded1 protein is sequestered with translation-inactive mRNAs in cellular foci when the translation conditions are altered (53,57,58). We reasoned that if polypeptide import into the ER was transiently blocked then Ded1 would form foci associated with the ER. We used temperature-sensitive (ts) mutants of the Sec61 and Sec62 proteins that form the translocon pore in the ER for the import of SRP-dependent polypeptides during translation (59,60). At the non-permissive temperature, these mutants block or disrupt the Sec61 channel and ER-associated translation is terminated. As a marker, we used the integrated red-fluorescent-tagged amino-terminal domain of Kar2 fused to the HDEL ER retention signal (YIPlac204TKC-DsRed-Express2-HDEL; Addgene, Watertown, MA) in the two *sec* strains; Kar2 is an ATPase that functions as a protein chaperone for refolding proteins within the lumen of the ER [(61) and reference therein], and consequently the Kar2 chimera serves as a marker of the ER lumen. We also used an ATPase-inactive Ded1-E307Q mutant (Ded1-DQAD) that has a high propensity to form cellular foci with sequestered mRNAs that are no longer undergoing translation.

We first looked at the distribution of proteins under permissive conditions (Figure 7A). The distribution of Kar2-RFP around the nuclear envelope (central cisternal ER), as interconnected tubules (tubular ER) and as a cortical halo inside the plasma membrane of the cell wall (PM-associated ER) was consistent with the locations of the ER in yeast; actively translating ribosomes are associated with all these ERs (62). Ded1-DQAD-GFP was uniformly distributed in the cytoplasm, largely excluded from the nucleus, and it formed occasional foci that were distributed at various positions in the cytoplasm (Figure 7A). Both the *sec61-ts* and *sec62-ts* strains showed equivalent phenotypes. The intensity of the fluorescence signals of both Kar2-RFP and Ded1-DQAD-GFP was highly variable between cells, which probably reflected different levels of protein expression between cells.

**Figure 7.**
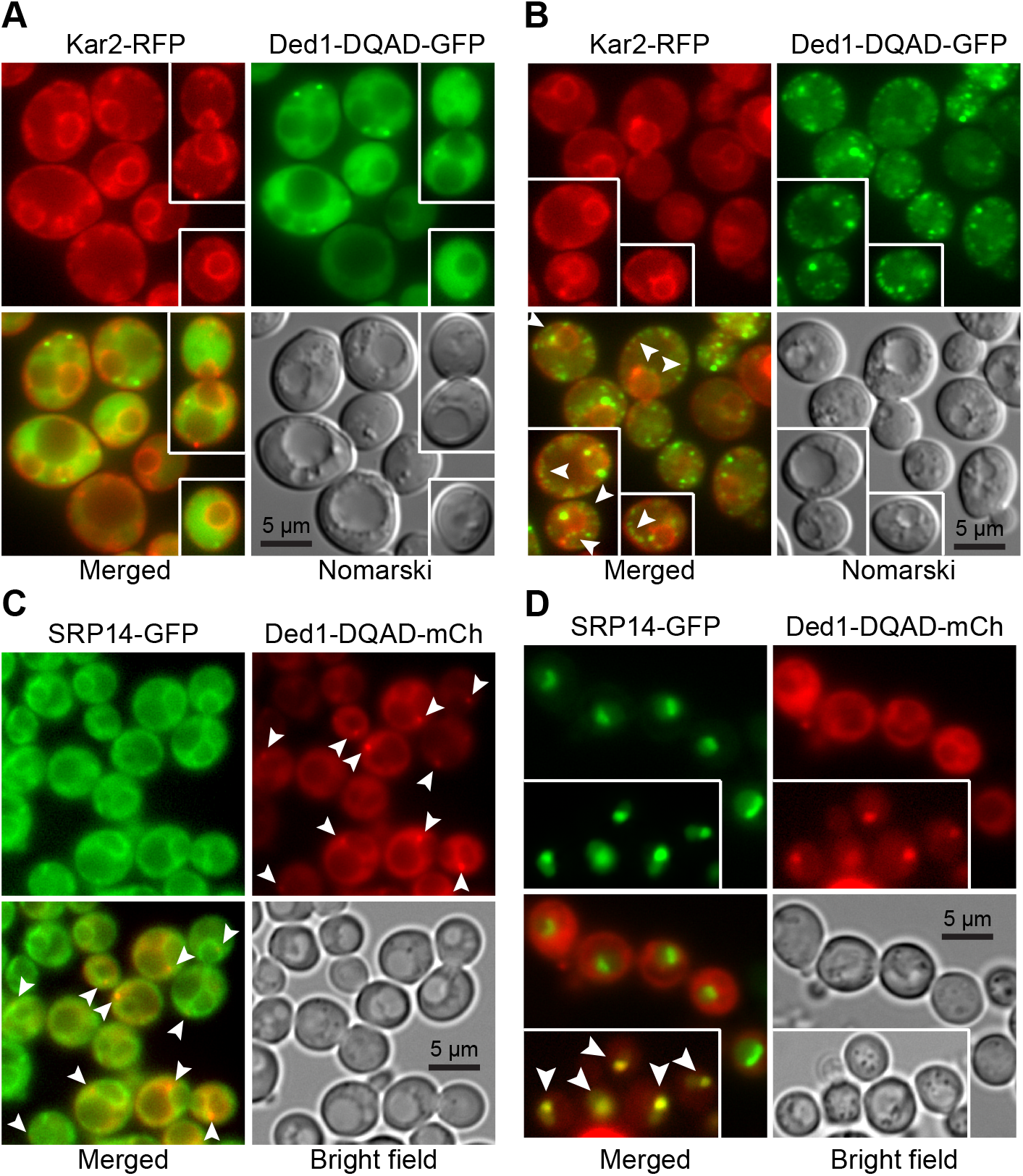
Cellular location of Ded1 relative to the ER and SRP proteins. (**A**) Ded1-DQAD-GFP was expressed in the *sec62* temperature-sensitive mutant with the integrated *KAR2-RFP* plasmid and grown to an OD_600_ of 1.0 at 24°C. (**B**) The same cells as in A were incubated for 15 min at the non-permissive temperature of 37°C prior to visualization. The arrowheads indicate positions where chains of Ded1-DQAD-GFP foci co-localized or co-associated with Kar2-RFP. (**C**) SRP14-GFP expressed from the chromosome and Ded1-DQAD-mCh expressed off the p415 plasmid were grown to an OD_600_ of 0.95 at 30°C. (**D**) SRP-GFP was overexpressed off the p413-PL plasmid and Ded1-DQAD-mCh was overexpressed off the p416-PL plasmid until an OD_600_ of 0.4 at 30°C in the *xpoI-T539C* yeast strain. Cells in the insert were treated with 10 µg/µl (∼200 nM) of leptomycin b for 1 h.

At 37°C, Kar2-RFP showed a similar cellular distribution as at 24°C for both *sec61-ts* and *sec62-ts* mutants, although it showed an increased frequency of aggregates with the ER (Figure 7B). In contrast, Ded1-DQAD-GFP showed a pronounced increase in the number of foci that were highly variable in size (Figure 7B). Many of these foci were closely associated with Kar2-RFP, particularly as a chain of foci on the cytoplasmic side of the ER around the plasma membrane, where the PM-associated ER was expected to be located, and as a chain of foci corresponding to tubular ER (arrowheads, Figure 7B). Both the *sec62-ts* and *sec61-ts* strains showed similar phenotypes. In some cases, the Kar2-RFP aggregates and Ded1-DQAD-GFP foci were near each other. Thus, Ded1 was associated with mRNAs that were no longer undergoing translation in close proximity to the ER at the non-permissive temperature. This result was consistent with previous work that showed that Ded1 is recovered with membrane-associated ribosomal-protein complexes (63).

We next asked if the SRP proteins showed similar properties. We used GFP-tagged SRP14 and SRP21 proteins that were expressed off the chromosome (GFP bank, Thermo Fisher Scientific, Waltham, MA) and the plasmid-encoded Ded1-DQAD-mCh mutant. The SRP14-GFP showed a weak but uniform signal in all the cells, where the protein was concentrated on the ER (Figure 7C). SRP21-GFP showed a similar profile (data not shown). In contrast, the plasmid-expressed Ded1-DQAD-mCh showed highly variable expression. We used cells grown under wildtype conditions or depleted for glucose, which promoted the formation of cellular foci. However, depending on the cellular growth we obtained a significant number of foci associated with the ER even under wildtype growth (arrowheads, Figure 7C). Thus, Ded1 was in close proximity to both the ER and the SRP proteins in the cell.

### Overexpressed SRP proteins accumulated in the nucleus and nucleolus

The biogenesis and metabolic pathway of the SRP RNP is complex, and it involves a large number of different steps [reviewed by (32,64,65)]. In yeast, the SRP proteins SRP14, SRP21, SRP68 and SRP72 are assembled on the SCR1 RNA probably in the nucleolus. Sec65 is in the nucleus, but there is some ambiguity about whether it accumulates in the nucleolus as well, although the equivalent mammalian SRP19 protein is found there (32,45,66). The partially assembled SRP complex is then exported to the cytoplasm through the XpoI/CrmI nuclear pore complex whereupon it binds with SRP54, which subsequently associates with the signal sequence of the partially translated polypeptide and causes the SRP to assemble on the 80S ribosomes. We previously showed that Ded1 actively shuttles between the nucleus and cytoplasm using the XpoI and Mex67 nuclear pores (13). Thus, it was possible that Ded1 associated very early with the SRP complex within the nucleus, and that it was important for the biogenesis or export of the complex.

The XpoI nuclear pore is known to export multiple cargoes, including ribosomal subunits, certain small nuclear RNAs, some viral RNAs and the assembled SRP complex (45,66–68). We used a yeast strain with a mutant *xpoI* allele that is sensitive to the bacterial toxin leptomycin b from *Streptomyces* to test whether Ded1 was involved in the XpoI-dependent export of the SRP (69); this strain contains a single mutation (XpoI-T539C) that makes the yeast protein sensitive to the drug (70). We previously showed that both the Mex67 and XpoI nuclear pore complexes must be disrupted to see a significant accumulation of Ded1 in the nucleus (13). We transformed this strain with plasmids expressing Ded1-mCh and with plasmids expressing either SRP14-GFP or SRP21-GFP, and we determined the locations of the tagged proteins.

The plasmid-encoded SRP14-GFP had highly variable expression between cells, but it showed a strong nuclear location that was often concentrated in crescent-shaped regions even in the absence of leptomycin b (Figure 7D, Figure 8A & 8B). In contrast, SRP21-GFP showed a diffuse location throughout the nucleus, which indicated that the overexpressed protein was not able to assemble or accumulate in the nucleolus (Figure 8C & 8D). However, it occasionally formed nuclear foci (arrowheads, insert Figure 8C). The expression of the plasmid-encoded Ded1-DQAD-mCh was likewise highly variable, and it was largely excluded from the nucleus even in the presence of leptomycin b (Figure 7D, Figure 8). However, in some instances with leptomycin b, where Ded1-DQAD-mCh was lightly expressed, we found that the protein accumulated in crescent-shaped regions with SRP14-GFP (arrowheads, insert Figure 7D). The Ded1-DQAD mutant binds RNA with a high affinity in the presence of ATP but it can not hydrolyze the ATP to recycle the complex. Thus, Ded1 could co-localize with the SRP complex in the nucleolus but only under conditions where Ded1 export was blocked. The absence of Ded1-DQAD in the nucleus and crescents in the absence of leptomycin argued that Ded1 was not needed for SRP assembly and export, but the data could not rule out this possibility.

**Figure 8.**
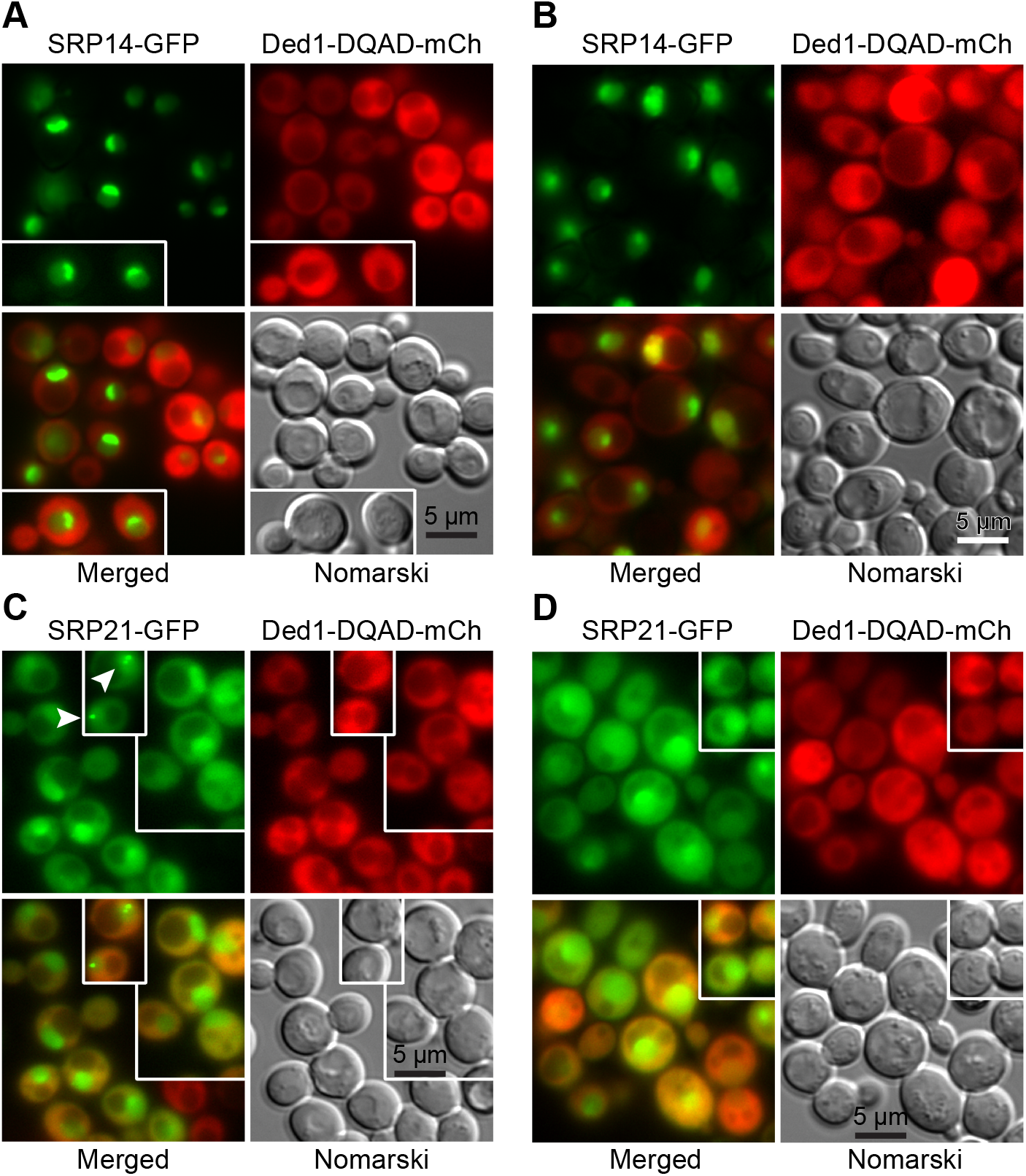
Over-expressed SRP14 and SRP21 accumulate in the nucleus. (**A**) SRP14-GFP was expressed off the p413 plasmid and Ded1-DQAD-mCh was expressed off the p416 plasmid in the *xpoI-T539C* yeast strain (70) and grown to an OD_600_ of 0.45 at 30°C. (**B**) The same as in **A** except that the cells were incubated for 60 min in the presence of 10 µg/ml of leptomycin b. (**C**) Same as A but with cells expressing SRP21-GFP. (**D**) Same as C but with cells incubated for 60 min with leptomycin b.

### Ded1 physically interacted with SRP factors

Our data indicated that Ded1 could bind SCR1, associate with the SRP complex and co-localize with the ER. Moreover, we obtained a genetic link between Ded1 and SRP proteins. However, it was unclear whether Ded1 physically interacted with the SRP proteins or indirectly through the SCR1 RNA (Figure 1). The metazoan SRP14 and SRP9 are known to bind 7SL in the Alu domain, SRP54 and SRP19 bind helices 6 and 8, and SRP68 and SRP72 bind around the junction between helices 5e, 5f, 6, 7 and 8 (27,35). However, yeast SRP14 is thought to bind the Alu domain as a homodimer and the role of SRP21 to date is largely speculative, although it is considered a structural homolog of SRP9 (36,38). Yeast Sec65 serves a similar role as SRP19, but it is considerably larger (47,49,71). The SRP bound on the 80S ribosomes shows an extended structure where the S-domain interacts with the exit channel containing the signal peptide and the Alu domain interacts with the entry region of the mRNA (37,39,40). However, SCR1 contains two hinge regions (Figure 1); it was possible that the SCR1 RNA was folded upon itself in its free form and that this brought the different regions of the SRP in close proximity. Thus Ded1 might interact with multiple different SRP proteins.

To test this, we subcloned the genes encoding the different proteins into pET22 and pET19 plasmids and then purified the recombinant proteins expressed in *Escherichia coli* on nickel-agarose columns. We then incubated the purified individual proteins or combination therein with purified Ded1, recovered the complexes with Ded1-IgG-Protein-A-Sepharose beads and separated the recovered proteins by SDS-PAGE (Figure 9).

**Figure 9.**
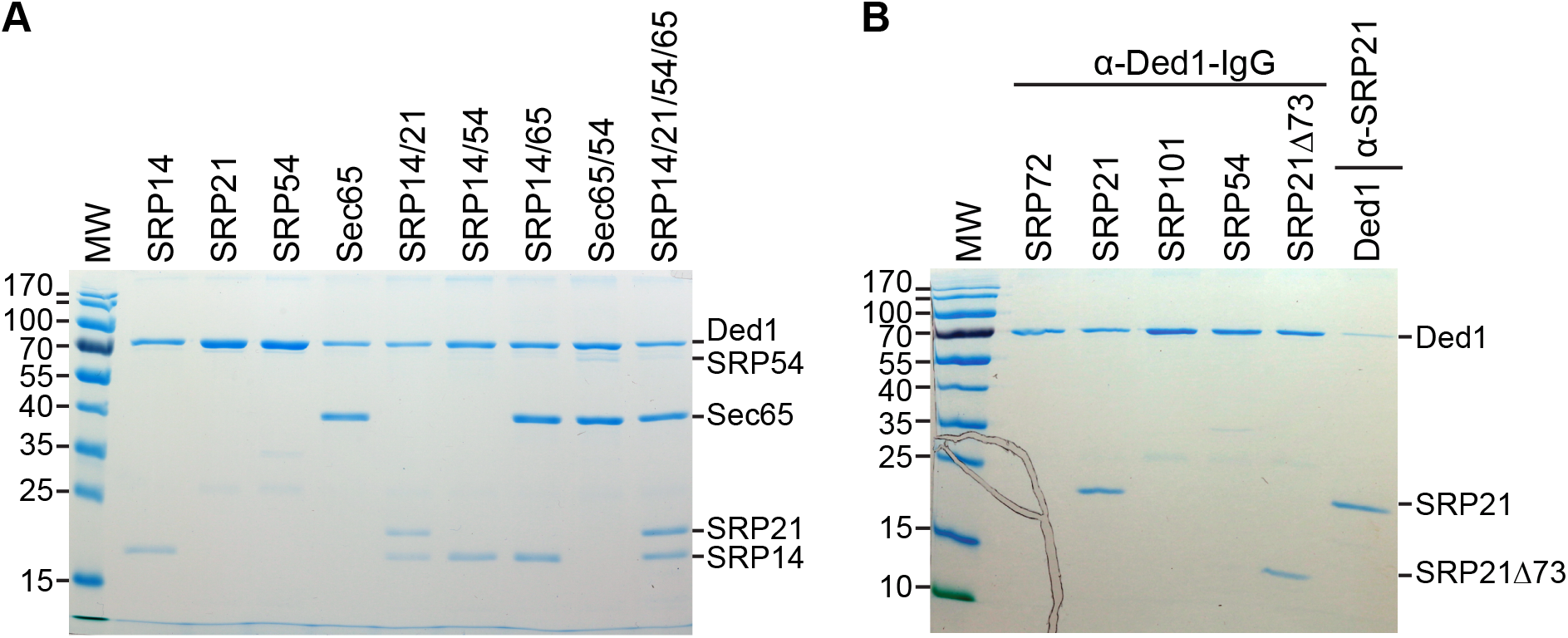
Ded1 physically interacted with the SRP proteins in the absence of RNA. (**A**) 4 µg of Ded1 was incubated with 4 µg of each SRP protein. The material was incubated 45 min at 30°C, immunoprecipitated with protein-A-Sepharose beads with Ded1-specific IgG, separated on a 12% SDS PAGE and visualized with coomassie blue. SRP68 and SRP72 migrated close to Ded1 and consequently were not unambiguously separated. (**B**) The same as A except 6 µg of the SRP proteins was used with 4 µg of Ded1. Proteins were recovered with Ded1- or SRP21-specific IgG as indicated.

The results showed that Ded1 formed stable interactions with SRP14 and Sec65. Unfortunately, SRP68, SRP72 and SRP101 migrated at positions on the SDS-PAGE that overlapped with Ded1; hence we were not able to obtain unambiguous results, but they appeared to have little or no affinity for Ded1. SRP54 did not stably associate with Ded1 by itself. In contrast, SRP21 was not consistently recovered with Ded1, which indicated that it formed weak interactions with the protein (Figure 9). However, SRP21 was consistently recovered with Ded1 in the presence of SRP14, which was consistent with SRP14 and SRP21 forming a stable heterocomplex as previously proposed (38). Likewise, SRP54 was recovered with Ded1 in the presence of Sec65; this was consistent with the two proteins being in close proximity on the S domain of SCR1 (47,49,71). Moreover, all four SRP proteins were recovered with Ded1 when incubated together.

The weak interactions between SRP21 and Ded1 were primarily through the amino-terminal domain because deleting the 73 carboxyl-terminal amino acids (SRP21Δ73) did not eliminate this affinity (Figure 9B). Finally, we recovered Ded1 in pull-down experiments with SRP21-specific IgG (Figure 9B). Thus, Ded1 was capable of forming protein-protein interactions with the SRP proteins in the absence of SCR1 RNA. Nevertheless, the presence of SCR1 RNA enhanced the recovery of all the proteins (see section: **Ded1 associated with SRP factors *in vivo***).

### SRP21 inhibited the SCR1-dependent ATPase activity of Ded1

Ded1 is an RNA-dependent ATPase. We previously showed that the nuclear and cytoplasmic cap-associated factors would stimulate the ATPase activity of Ded1 in the presence of RNA (13). We wondered whether the SRP proteins would alter the enzymatic activity of Ded1 as well and whether it would be preferential for the SCR1 RNA, which would be the authentic substrate for the assembly of the SRP proteins. We tested this with an *in vitro*, T7-polymerase-transcribed SCR1 RNA that was equivalent to the endogenous SCR1 except that the 5’ terminal nucleotide was replaced with a guanosine to facilitate transcription. As a control we used a fragment of the actin pre-mRNA precursor containing short exon sequences and the entire intron. In addition, the actin transcript was of similar size to SCR1 (605 nts and 552 nts, respectively).

Many of the SRP proteins at nearly a 30-fold excess over Ded1 inhibited the RNA-dependent ATPase activity of Ded1 somewhat for both actin and SCR1, although SRP54 seemed to enhance the activity slightly, especially with actin (Figure 10A). This was not unexpected because the SRP proteins were largely basic and positively charged under the reaction conditions (pH 7.5; Table 3); the proteins would be expected to nonspecifically associate with the RNAs and thereby reduce the effective concentration of the RNAs accessible to Ded1. SRP21 showed a much stronger inhibition, especially with SCR1, but it was the most basic (pK_i_ = 11.14) of the SRP proteins. To further elucidate the nature of the inhibition, we compared the ATPase activity of Ded1 with SCR1 and actin RNAs with SRP21. SRP21 is considered the structural homolog of SRP9, and in yeast it probably forms a complex with a homodimer of SRP14 that binds the Alu domain of SCR1 (36,38). Thus we tested to see if SRP14 would enhance the inhibitory effects of SRP21.

**Figure 10.**
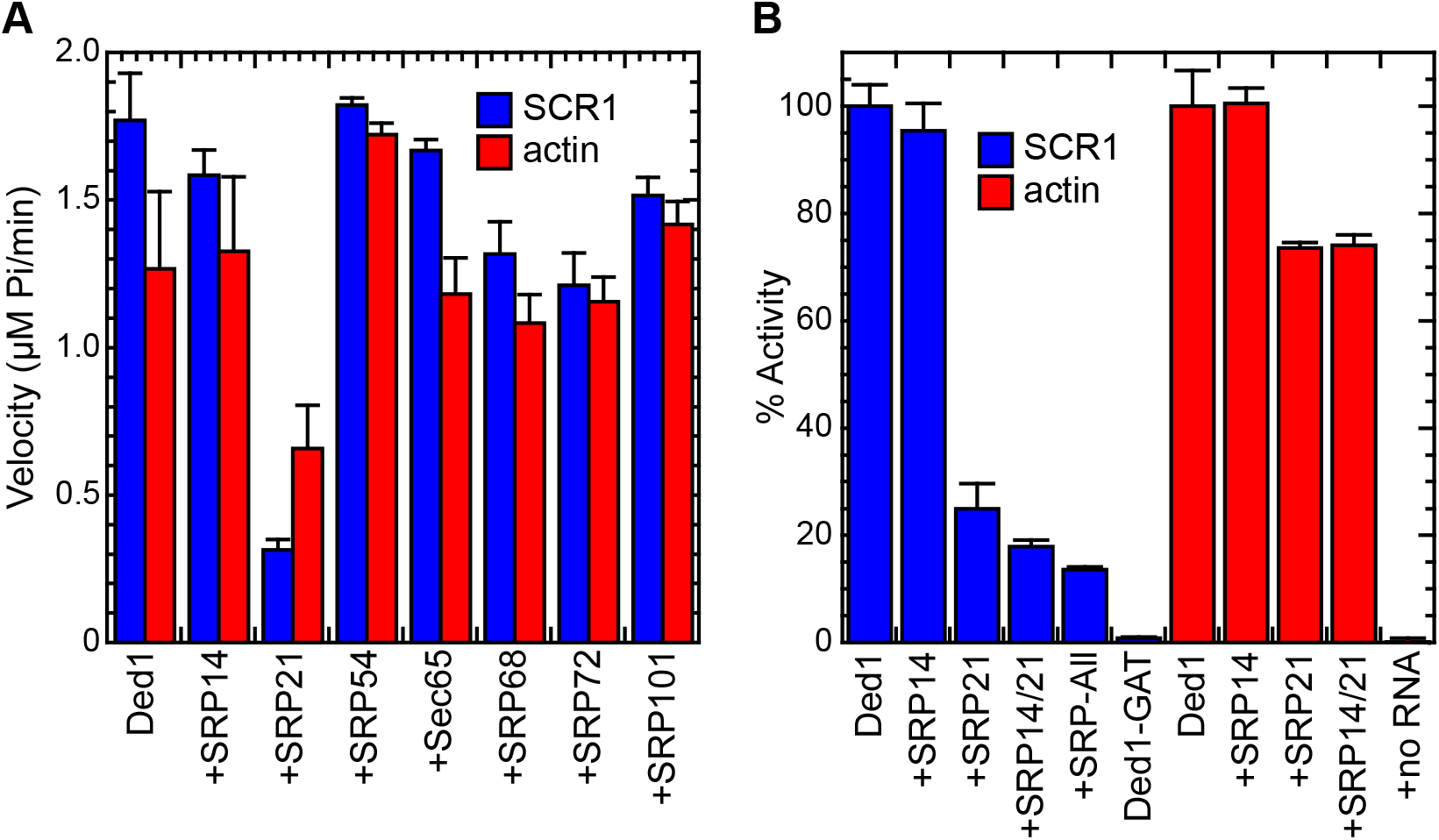
The SRP proteins inhibited the ATPase activity of Ded1. (**A**) Reactions were undertaken with 7 nM Ded1, 200 nM of the SRP proteins, 1 mM ATP and 23 nM of SCR1 or actin RNA. The reaction velocities were measured over 40 min at 30°C. (**B**) Reaction were done as in A but with 23 nM of the SRP proteins except SRP14, which was used at 46 nM to form the homodimer, and 23 nM RNAs. The reaction velocities were normalized relative to the activity of Ded1 in the presence of the RNA (SCR1 or actin) alone. +SRP-All, Ded1 was incubated with SRP14, SRP21, SRP54, Sec65, SRP68, and SRP72; Ded1-GAT, a Ded1 P-loop mutant that lacks ATPase activity; +no RNA, Ded1 was incubated in the absence of an RNA substrate with the SRP proteins. The mean and standard deviations are shown for two independent experiments in panel A and for three in panel B. The lower error bars were deleted for clarity.

Equimolar concentrations of both the actin precursor and SCR1 stimulated the ATPase activity of Ded1, but the stimulation was not equivalent. Moreover, there was variability in the stimulatory effects with different RNA preparations, which probably reflected variability in the folding of the RNAs during preparation. Indeed, others have shown that the smaller human 7SL RNA is difficult to recover as a homogeneous structure *in vitro* (41). Thus, to facilitate comparisons we normalized the activity relative to that of Ded1 with the RNA alone and used the same RNA preparations for comparisons. In addition, we used 8.5-fold less of the SRP proteins to emphasize the differences.

SRP14 may have slightly inhibited Ded1 with SCR1 but it had little affect with actin (Figure 10B). In contrast, SRP21 inhibited the ATPase activity of Ded1 with SCR1 by about 75% but only by 25% with actin. This indicated that there was some nonspecific inhibition, but that the strongest inhibition was obtained with the authentic substrate of SRP21. Addition of SRP14 reduced the activity of Ded1 in the presence of SRP21 by an additional 8% for SCR1 but showed no additional reduction with actin. Thus, SRP14 enhanced the SRP21-dependent inhibition of Ded1 with SCR1 RNA. Addition of the other SRP proteins to SCR1 further reduced the activity by about 4%. None of the purified SRP proteins showed any intrinsic ATPase activity in the absence of Ded1, and none of the SRP proteins affected the ATPase activity of Ded1 in the absence of RNA (Figure 10B and data not shown).

The data indicated that SRP21 in the presence of its authentic substrate was the most effective at inhibiting Ded1. Thus, it was likely that SRP21 bound to the Alu domain of SCR1 formed the most effective inhibitory structure. We tested this with deletions of the S domain (SCR1ΔS1) and Alu domain (SCR1ΔAlu). Previous work has shown that the folding of mammalian 7SL RNA is difficult, and it needs a temperature step for refolding involving slow cooling in the presence of monovalent cations; moreover, the assembly of the SRP proteins is complicated (41). The yeast SCR1 RNA is about two-fold larger with a number of additional hairpins, so we anticipated difficulties in obtaining a functional homogenous structure (Figure 1). Thus, we assayed various permutations of pre-incubating the RNA with the various proteins prior to adding the ATP, but they all yielded similar results. Ded1 was significantly less active with both SCR1ΔS1 and SCR1ΔAlu then full-length SCR1 at equimolar concentrations of RNA, but the RNAs were 77% and 58%, respectively, of the size of SCR1 (Figure 11A). This may account for the reduced ATPase activity, but it was possible that Ded1 was activated by specific structures within the SCR1 RNA that were absent or misfolded in the deletions. SRP21 inhibited the ATPase activity for SCR1 and to a lesser extent actin, but it had little inhibitory affect on the SCR1 deletions, which was consistent with a structure-dependent inhibition. We also tested a carboxyl-terminal deletion of SRP21 (SRP21Δ73) that lacked the amino acids that did not correspond to those of mammalian SRP9 (38); it showed significantly less inhibition, which was consistent with it playing a role in the SRP21 interactions with SCR1 and Ded1 (Figure 11A).

**Figure 11.**
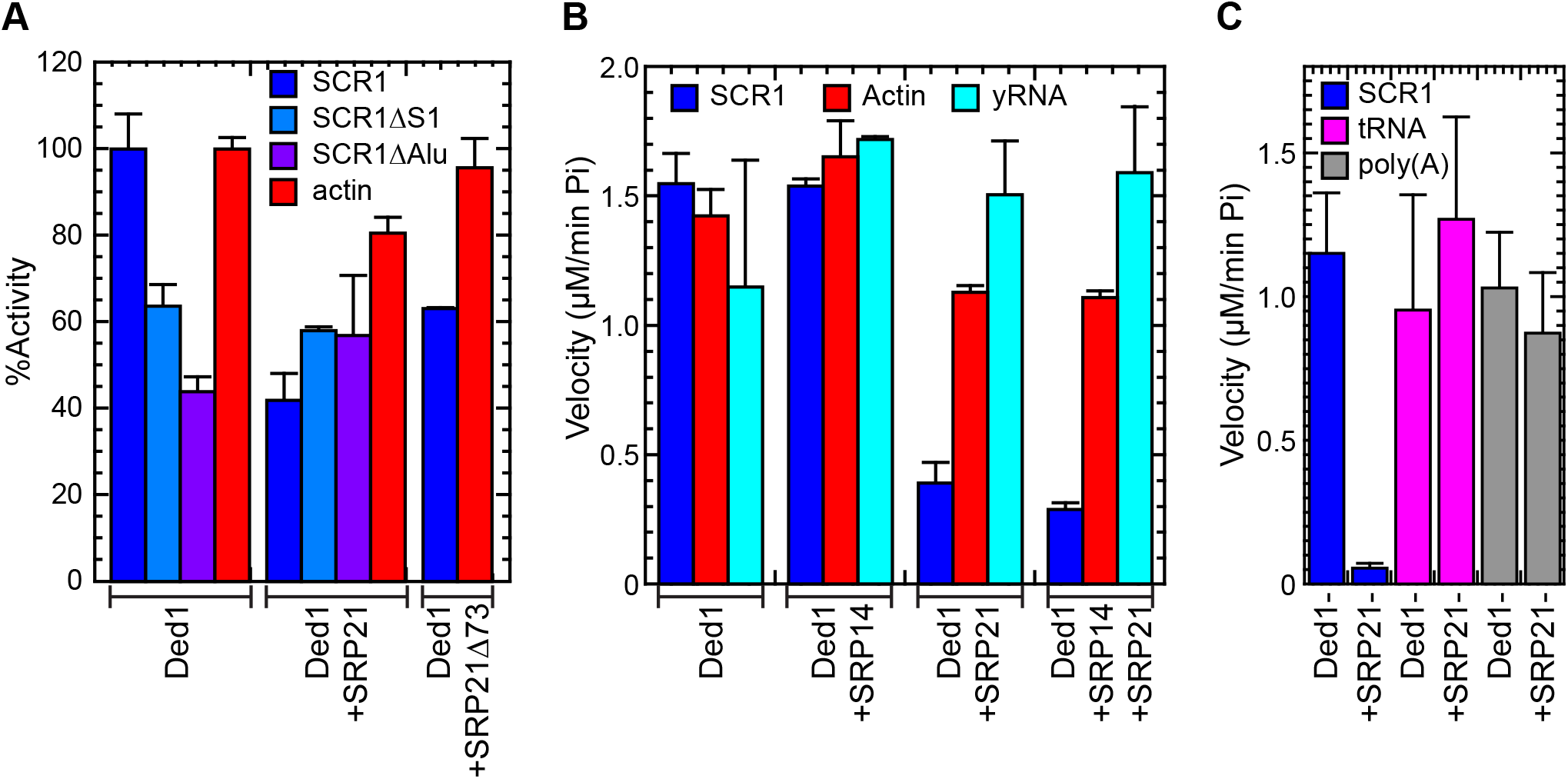
The RNA-dependent effects of SRP21 on the ATPase activity of Ded1. (**A**) Ded1 was pre-incubated with the RNAs at 30°C for 30 min. SRP21 or SRP21Δ73 were then added at 200 nM with 1 mM ATP, and the ATPase velocity was measured over 40 min. The mean and standard deviations are shown for two independent experiments. (**B**) Ded1 at 7 nM was incubated with 23 nM SCR1, 23 nM actin or 0.14 µg/µl yeast RNA and with 1 mM ATP. SRP21 was used at 23 nM and SRP14 at 46 nM (to form homodimer). The reaction velocities were measured over 40 min at 30°C. The mean and standard deviations are shown for three independent measurements are shown for SCR1 and actin and for two independent measurements for yeast RNA. (**C**) Reactions were done as in B. Ded1 at 7 nM was incubated with 23 nM SCR1 (equivalent to 0.0039 µg/µl) or with 0.12 µg/µl of tRNA or poly(A). The SRP21 was used at 200 nM. The mean and standard deviations are shown for three independent measurements. The lower error bars were deleted for clarity.

The ATPase activity of Ded1 is stimulated by various RNAs containing single-stranded regions, but it is most activated by poly(A)-containing RNAs (72). We repeated the ATPase assays with purified yeast RNA. We needed to use 0.12–0.14 µg/µl of yeast RNA to obtain similar levels of activation of Ded1 as 23 nM of SCR1 (0.0039 µg/µl) or actin (0.0044 µg/µl; Figure 11B). Thus, yeast RNA was ∼30-fold less effective at stimulating the activity. SRP21 and SRP14 showed no inhibitory effects, and they may have actually stimulated the ATPase activity of Ded1 somewhat, perhaps by acting as RNA chaperones to increase the accessibility of the RNA to Ded1. However, the yeast RNA was a heterogenous mix that may have had both activating and inhibitory RNAs. Thus, we repeated these experiments with yeast tRNAs and poly(A) RNA at 0.12 µg/µl (Figure 11C). Both RNAs needed ∼30–fold higher concentration than for SCR1 to stimulate the ATPase activity of Ded1 to similar levels, but SRP21 had little affect on the activities. Thus, Ded1 and SRP21 preferentially bind RNAs with certain sequences or structural features.

### SRP21 did not block Ded1 binding to SCR1

The previous results indicated that Ded1 was either blocked from binding the SCR1 RNA or that its ATPase activity was inhibited by protein-protein contacts with SRP21 bound on the RNA. To test this, we did electrophoretic mobility shift assays (EMSA) with the different proteins and RNAs. Our previous work showed strong, concentration-dependent binding of Ded1 to short oligonucleotides in the presence of AMP-PNP with a K_1/2_ of ∼40 nM and weak binding in the presence of ADP or in the absence of a nucleotide (73). We repeated these experiments with the longer RNAs, but we separated the products on agarose gels containing ethidium bromide. Similar results were obtained when the gels were run in the absence of ethidium bromide, which was then soaked into the gels after electrophoresis (data not shown).

A 5- to 10-fold excess of Ded1 was able to displace the majority of both SCR1 and actin (Figure 12A). SCR1 typically migrated as a distinct band but actin often showed more heterogeneity, which probably reflected more profound conformational heterogeneity. This varied somewhat between RNA preparations. In contrast, SRP21 preferentially bound SCR1 RNA over actin (Figure 12B). Moreover, it seemed to form large molecular-weight aggregates that only partially migrated into the gels. Deleting the carboxyl-terminal sequences of SRP21 (SRP21Δ73) largely eliminated the binding affinity, indicating that these sequences were either important for binding or for maintaining the correct conformation of the protein (Figure 12C).

**Figure 12.**
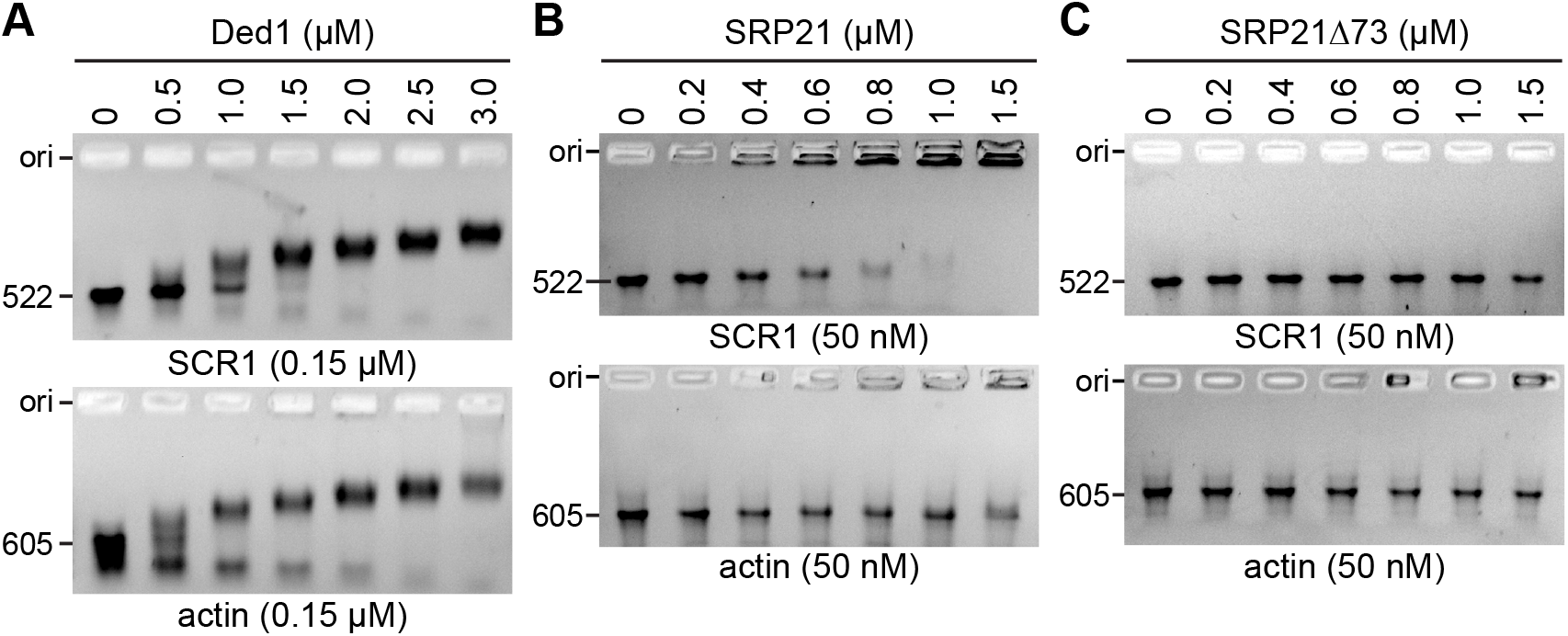
RNA binding assays of Ded1 and SRP21. **(A**) Ded1 binds SCR1 and actin with similar affinities. The indicated quantities of the Ded1 protein was incubated with 0.15 µM of the indicated RNAs and then separated on a 1% agarose gel in the presence of ethidium bromide. Ori, loading well of agarose gel. (**B**) The indicated quantities of the SRP21 proteins were incubated with 50 nM of the indicated RNAs and separated on a 1% agarose gel. (**C**) SRP21 deleted for the 73 carboxyl-terminal residues that are not structurally conserved in mammalian SRP9.

We next asked what affect SRP21 would have on Ded1 binding. The results showed that SRP21 had little affect on Ded1 binding, but the retarded bands tended to migrate as higher molecular-weight complexes in the presence of SRP21 for SCR1 (Figure 13A). We previously showed that the carboxyl-terminal domains of DEAD-box proteins, including Ded1, are important for high affinity binding to RNAs (54). Consistent with this, deleting 78 amino acids from the carboxyl terminus of Ded1 largely eliminated RNA binding (Figure 13B). However, addition of SRP21 had little affect on Ded1 binding even though a small amount of material was sequestered near the origin of the gel (Figure 13C). Instead, Ded1 seemed to actually reduce the binding affinity of SRP21 for the RNA, and this was true for the carboxyl-terminal deletion of Ded1 as well (Figure 13C). Thus, although SRP21 modulated the ATPase activity of Ded1, Ded1 seemed to modulate SRP21 binding, perhaps through interactions with the amino-terminal domain of Ded1 or the RecA-like core.

**Figure 13.**
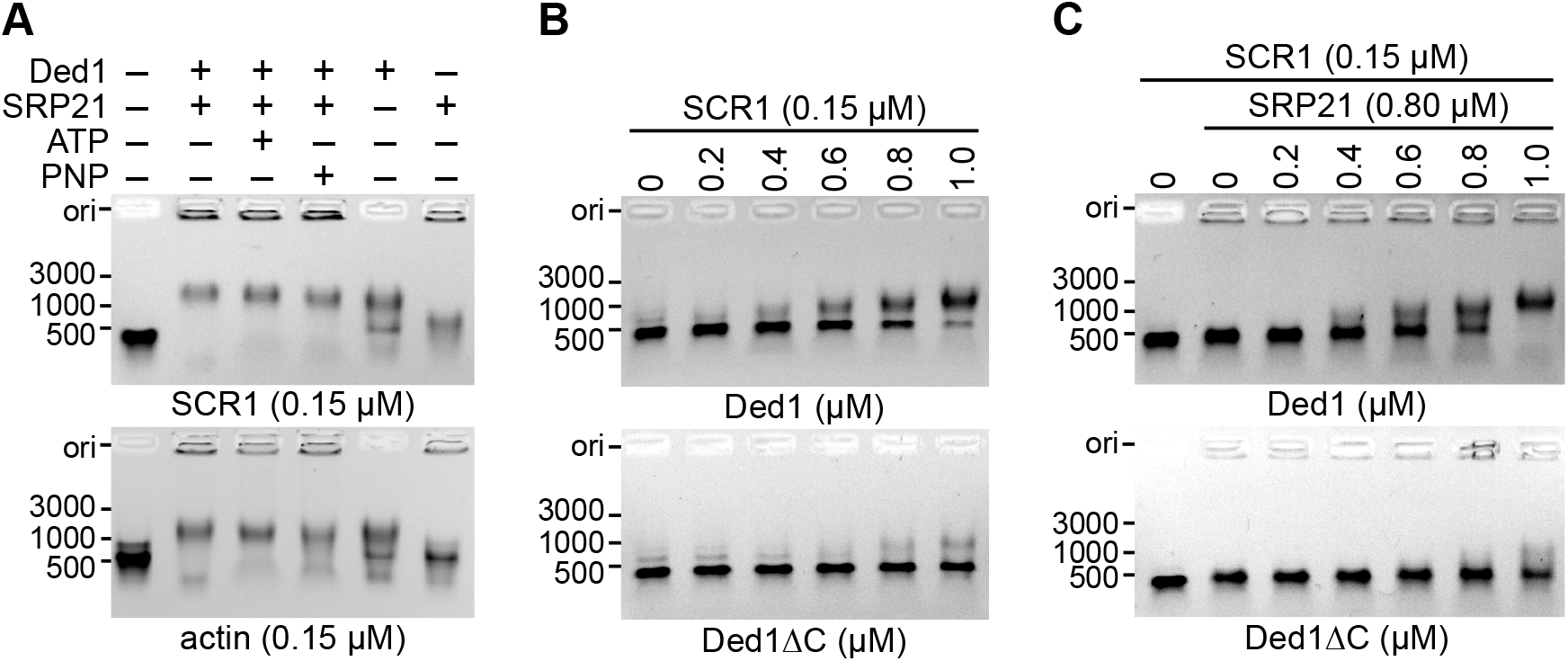
Electrophoretic mobility shift assays of Ded1 with SCR1 (522 nts) and actin (605 nts) RNAs. Proteins were incubated with the RNA and separated under nondenaturing conditions on 1% agarose gels containing ethidium bromide. The markers indicate the positions of the major bands of the GeneRuler DNA ladder (Thermo Scientific). Ori, loading well of agarose gel. (**A**) Ded1 (0.8 µM) and SRP21 (0.8 µM) were incubated with 0.15 µM of either SCR1 or actin RNA in the presence or absence of 5 mM ATP or AMP-PNP (PNP). (**B**) Increasing concentrations (in µM) of Ded1 or an 78 amino-acid, carboxyl-terminal deletion of Ded1 [Ded1ΔC; (54)] was incubated with 0.15 µM SCR1 RNA and 5 mM AMP-PNP. (**C**) Increasing concentrations of Ded1 was incubated with 0.15 µM SCR1 RNA, 0.80 µM SRP21 and 5 mM AMP-PNP.

Finally, we asked what structural features of SCR1 were recognized by the proteins. These experiments were more ambiguous because the binding site of SRP21 is largely unknown and because there was no guarantee that the RNAs deletions would fold into the anticipated conformations. The results showed that both Ded1 and SRP21 were able to bind the SCR1 RNAs deleted for the Alu and S domains (Figure 14). Thus, SRP21 probably recognized multiple features of the SCR1 RNA. Moreover, the strong binding of Ded1 to the deleted SCR1 RNAs did not correlate with the reduced ATPase activities of Ded1 with these RNAs (Figure 11A). Therefore, either Ded1 could bind the RNAs in a nonproductive form or the ATPase activity of Ded1 was modulated by the different structures.

**Figure 14.**
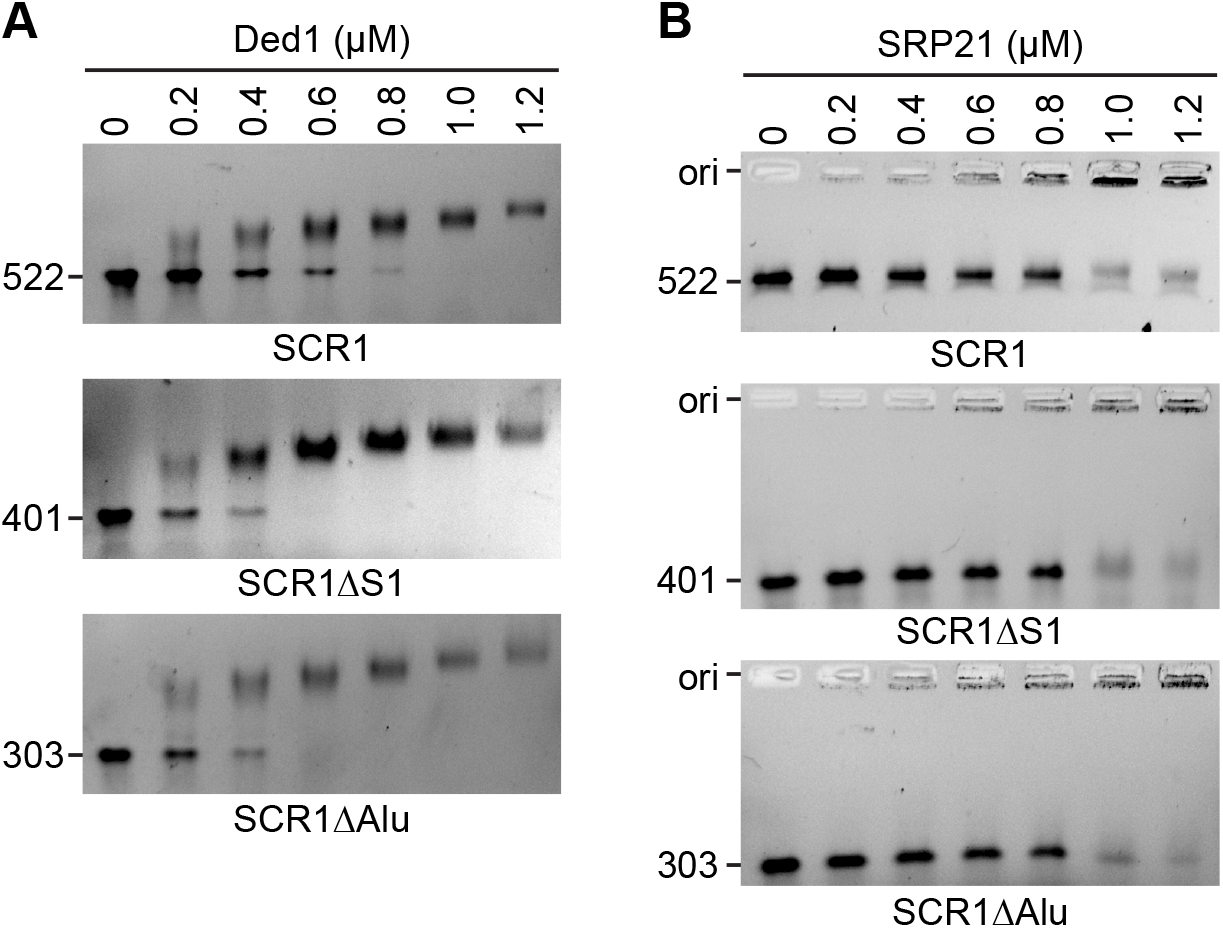
Ded1 and SRP21 bind various regions of SCR1. The indicated quantities of protein were incubated with the indicated RNAs and separated on a 1% agarose gel in the presence of ethidium bromide. (**A**) The indicated concentrations of Ded1 (in µM) were added to 0.15 µM of the different SCR1 RNAs in the presence of 5 mM AMP-PNP. (**B**) The indicated concentrations of SRP21 (in µM) were added to 0.15 µM of the different SCR1 RNAs. Ori, loading well of agarose gel.

## DISCUSSION

Our experiments show that Ded1 is an SRP-associated factor. It physically interacts with the SCR1 RNA and many of the SRP proteins both *in vitro* and *in vivo*. It is genetically linked to these proteins, and it co-sediments with the SRP factors in sucrose gradients. The RNA-dependent ATPase activity of Ded1 is inhibited by SRP21 and this inhibition is much more pronounced in the presence of SCR1 RNA, the authentic substrate of SRP21. Although there is probably conformational heterogeneity of the RNAs, SRP21 preferentially binds SCR1 RNA over actin RNA, which indicates that it contains or forms the necessary elements for high-affinity SRP21 binding. Likewise, the ATPase activity of Ded1 is preferentially activated by SCR1 and actin RNAs over an equivalent concentration whole yeast RNA, tRNA or poly(A) RNA, which indicates that it is recognizing specific features or structures of these RNAs. The nature of these features or structures is unclear. Finally, Ded1 co-localizes in cellular foci with the ER-associated mRNAs, and it occasionally co-localizes with SRP proteins in the nucleolus.

The role of SRP21 to date is largely unclear. It is considered as the structural homolog of metazoan SRP9, which forms a heterodimer with SRP14 on the Alu domains of 7SL RNA, even though there is little or no sequence homology (38). The amino-terminal residues of SRP21 are capable of forming similar structural features as SRP9, but it is over 80% bigger; the carboxyl-terminal sequences are thought to compensate for the abbreviated Alu domain of yeast SCR1, which lacks the characteristic hairpins H3 and H4 (38). Moreover, yeast SCR1 is about 75% bigger than metazoan 7SL, and it contains additional structures between the Alu and S domains (33,34). SRP21 may be needed to stabilize or form the correct conformation of the SCR1 RNA, and thus it may need to recognize multiple structures or features of the RNA. Consistent with this, SRP21 binds full-length SCR1 RNA and the deletions *in vitro* with similar affinity. In contrast, it has weak affinity for the actin RNA. The carboxyl-terminal sequences of SRP21 are important for this affinity. This is consistent with SRP21 forming a complex with the SRP14 homodimer as previously proposed (38).

SRP21 inhibits the RNA-dependent ATPase activity of Ded1, but it is much more effective in the presence of SCR1 RNA than actin RNA. In contrast, Ded1 binds SCR1 and actin RNAs with similar affinities and it is activated to similar extents. Under these circumstances, one would expect SRP21 to reduce Ded1 binding to SCR1 but not to actin because SRP21 would reduce the number of potential binding sites for Ded1 on SCR1. But this is not the case, and if anything SRP21 seems to enhance Ded1 binding to SCR1 slightly. The inhibition is due to protein-protein contacts, but SRP21 is less stably associated with Ded1 in the absence of SRP14 or SCR1 RNA. Thus, SRP21 probably forms a specific inhibitory structure with Ded1 in the presence of SCR1 RNA. This is consistent with a functional regulation of the ATPase activity of Ded1 in the context of the SRP complex.

Ded1 is an ATP-dependent RNA binding protein, and it is capable of forming long-lived complexes with RNA in the presence of a nonhydrolyzable analog of ATP *in vitro* (8). Ded1 is considered a translation-initiation factor [(14,15) and references therein], but crosslinking studies on DDX3 show most of the interactions on the open reading frames of a subset of the mRNAs (74). We obtained similar results with Ded1 (data not shown). Thus, Ded1 remains associated with the ribosomes during translation elongation, which can be seen in polysome profiles as well [this work and (13)]. Ded1 likewise is found with membrane-associated ribosomes (63). Thus, Ded1 may play important roles in translation elongation as well as in initiation—including membrane-associated translation.

Ded1 is associated with 90S ribosomal precursors, which may indicate a role of Ded1 in SRP assembly in the nucleolus (75). We do see occasional co-localization of the Ded1-DQAD mutant with overexpressed SRP14 in crescent-shaped structures in the nucleus that are consistent with this possibility, but SRP21 has a diffuse location within the nucleus, and it is never seen concentrated in the crescent-shaped structures. Thus, it may associate with the SRP complex outside the nucleolus or it may be transiently located within the nucleolus, as has been proposed for Sec65 (45). Thus, we can not rule out a role for Ded1 in the biogenesis of the SRP complex in the nucleus that is regulated by SRP21. Under these circumstances, Ded1 may associate early with the SRP complex and remain attached even when the complex binds the 80S ribosomes. This would provide a possible mechanism by which ER-specific mRNAs are selected for translocation on the ER. Interestingly, DDX3 crosslinks to 7SL RNA as well (74). Thus, although metazoans lack a clear equivalent to SRP21, DDX3 may also be intimately connected to SRP-dependent translation.

On the basis of these observations, we propose the following model for the role of Ded1 (and DDX3-like proteins) in membrane-associated translation. Ded1 interacts with cap-associated factors and with Pab2 bound on the 3’ poly(A) tail of the mRNA (not shown). The 3’ UTR is considered important for SRP-dependent targeting of mRNAs [reviewed by (76)]; but the SRP is not known to directly interact with the mRNA, and it may interact through another RNA binding protein (77). Ded1 (and DDX3) could serve this role as it interacts with both 5’ and 3’ components of the mRNA [(13,16) and references therein]. Ded1 remains attached to the mRNA during scanning by the 43S ribosomes, assembly of the 48S ribosomes and eventual formation of the 80S ribosomes at the AUG start codon (Figure 15A). This is consistent with crosslinking experiments of ribosomal RNA that show Ded1 near the mRNA entry channel (43). The RNA-dependent ATPase activity of Ded1 is uninhibited, and it is able to translocate on the mRNA with the ribosomes through rapid cycling between the “open” and “closed” conformations, and it may further stabilize the ribosome-mRNA complex during scanning, assembly and translation (14). During this time, an inactive form of the SRP may associate with the mRNA-ribosome-Ded1 complex through postulated 3’ UTR factors that are specific for SRP-dependent translation while the ribosomes are still part of the soluble fraction during a pioneering round of translation as previously proposed (77). The Alu domain of the SRP would be easily displaced from the ribosomes by elongation factors (78). Ded1 may play an important role in the assembly and stabilization of this complex because it can interact with all the relevant factors. Alternatively, Ded1 may help associate the SRP on the ribosomes once the signal peptide is sufficiently long according to the classical model (79).

**Figure 15.**
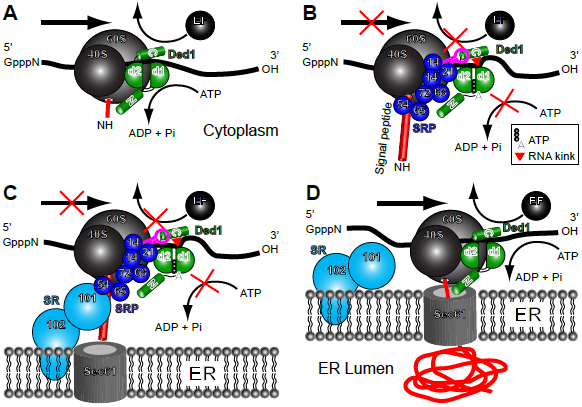
Model for the role of Ded1 in SRP-dependent translation. (**A**) Ded1 (shown in green) associates with the mRNA during translation initiation and remains attached to the mRNA in front of the ribosomes. It consists of RecA-like domains 1 (d1) and 2 (d2), an amino-terminal domain (N) and a carboxyl-terminal domain (C). The RNA-dependent ATPase activity of Ded1 is unaltered, and it is often in the “open” conformation with weak affinity for the RNA; it is able to translocate with the ribosomes during translation. (**B**) The SRP (shown in blue) associates with ribosomes translating mRNAs (or undergoes conformational changes in the case of pre-bound SRP) when the signal peptide leaves the exit channel and obtains a certain length. Ded1 may help in assembling and stabilizing the complex. Conformational changes of the SRP causes SRP14 to block the entry channel and prevent the eEF2 elongation factor (EF) from binding the ribosomes, which pauses elongation. Ded1 may bind part of the Alu domain of SCR1, shown in magenta, during these conformational changes to promote SRP14 binding to the ribosomes. At the same time, SRP21 inhibits the ATPase activity of Ded1, which forms the “closed” conformation with high affinity for the RNA. This ATP-bound form of Ded1 kinks the RNA (red triangle) on domain 1 and locks Ded1 on the RNA. This prevents the ribosomes from frame shifting (sliding) on the RNA and perhaps stabilizes the ribosome-mRNA complex to prevent premature termination of translation. (**C**) The paused mRNA-ribosome complex associates with SRP receptor (SR) factors SRP101 and subsequently SRP102, which brings the mRNA-ribosome complex to the Sec61 ER translocon. (**D**) The SRP complex dissociates from the ribosomes, the ATPase activity of Ded1 is restored and translation continues. Note that this model also applies to the SRP-dependent import of polypeptides with internal transmembrane domains, and it does not preclude the possibility that multiple Ded1 molecules are involved, that the SRP associates multiple times with the ribosomes during elongation or that the SRP-associated ribosomes remain on the ER over multiple rounds of translation.

In the next step, the signal peptide binds in the hydrophobic groove of the GTPase SRP54 that subsequently causes conformational changes of the SRP and its interactions with the ribosome [(40,78,80) and references therein]. SRP14 bound on the Alu domain of SCR1 binds at the GTPase center located at the 40S-60S interface and thereby transiently blocks the GTPase elongation factor eEF2 from binding (37,39,40). At the same time, SRP21 binds to Ded1 and inactivates its ATPase activity. This results in Ded1 maintaining a closed conformation that has high affinity for the RNA but that also crimps the RNA bound on RecA domain 1; this prevents both Ded1 and the ribosomes from sliding on the mRNA by effectively clamping the mRNA (Figure 15B). Another factor than SRP9 may play this role in metazoans. The SRP complex undergoes conformational changes during this time, and Ded1 may also bind the SCR1 RNA through its carboxyl terminus to facilitate the subsequent interactions of SRP14 with the ribosomes.

The absence of DDX3-like RNA helicases in *in vitro* reconstituted systems might explain why this pausing is often short or absent (81). Ded1 may also stabilize the paused ribosomes to prevent premature termination or frameshifting. The paused ribosomes then associate with the peripheral-membrane GTPase SRP101 and the integral-membrane protein SRP102 that form the SRP receptor (SR) complex (Figure 15C). Once associated with the Sec61 translocon, the SRP complex undergoes further conformational changes and is either released from the ribosome or assumes an inactivated form on the ribosomes. The ATPase activity of Ded1 is restored and translation can resume (Figure 15D).

Ribosome pausing events are important for other translational events in addition to SRP-dependent protein translocation. For example, pausing is associated with co-translational protein folding, protein targeting, mRNA and protein quality control, and with co-translational mRNA decay [reviewed by (82)]. Ded1 (and DDX3) may be intimately associated with these events by a similar mechanism but with other associated factors besides the SRP proteins. Likewise, ribosome pausing is associated with frameshifting events [reviewed by (83,84)]. Ded1 is important for L-A RNA virus replication (85), and it undergoes a -1 frameshifting event during translation of the gag-pol gene [reviewed by (86)]. Similarly, retroviruses, such as HIV-1, undergo a −1 frameshifting event during translation [reviewed by (87)]. DDX3 is important for HIV-I replication, and the virions of retroviruses in general are enriched in 7SL RNA (23,88). Thus, the Ded1/DDX3 subfamily of proteins may play central roles in gene expression by regulating not only translation initiation but translation elongation as well.

Finally, we note that the bacterial polypeptide-translocase SecA is a superfamily 2 “RNA helicase” that has a RecA-like core structure that is very similar to the DEAD-box proteins (89). It is intimately associated with the SecYEG translocon, and it uses ATP to drive post-translational polypeptides through the pore into the periplasm [reviewed by (26)]. Recent work has shown that SecA mimics the properties of the SRP (90,91). Thus, the use of superfamily 2 proteins for polypeptide translocation across membranes may be conserved throughout evolution. The biological roles and substrates of these proteins may not be limited to nucleic acids.

## MATERIAL AND METHODS

### Constructs

The pMW295 and pMW299 plasmids encoding the SRP proteins and SCR1 were a kind gift from Martin R. Pool (44). They were used as templates to PCR-amplify the individual genes and SCR1 RNA. Other constructs are as described below. The oligonucleotides used and cloning strategies are shown in more detail in Supplementary Table 1. The SRP proteins were PCR amplified with the corresponding SRP _up and _low oligonucleotides off the pMW295 and pMW299 plasmids (44). The PCR products were digested with SpeI and XhoI, gel purified, and cloned into the equivalent sites of the yeast plasmids 2HA-p415 and 2HA-p424 with *ADH* promoters and *LEU2* and *TRP1* markers, respectively (92). Except for SRP68 and Sec65, all constructs were subcloned into the NdeI and XhoI sites of pET22b.

Because of internal NdeI sites, the pET19b versions of SRP68 and Sec65 were amplified with additional oligonucleotides. SRP68 was PCR amplified with SRP68_up2 and SRP68_low2. Sec65 was PCR amplified with Sec65-pET_up and Sec65-pET_low. The PCR products were digested with XhoI and BamHI, gel purified and cloned into the equivalent sites of pET19b.

SRP101 and SRP102 were amplified off purified chromosomal DNA using oligonucleotides with BamHI and XhoI sites because of an internal SpeI site in SRP101. SRP101 was PCR amplified with SRP101_up and SRP101_low. SRP102 was PCR amplified with SRP102_up and SRP102_low. The PCR products were digested with BamHI and XhoI, gel purified and cloned into the equivalent sites of 2HA-p424. The constructs were subcloned into the NdeI and XhoI sites of pET22b.

The SRP14 and SRP21 wildtype and carboxyl-terminal deletions were cloned into the p413 plasmid containing a *HIS3* marker and *ADH* promoter by using an oligonucleotide complementary to the amino-terminal HA tag of the previous p415-PL-HA constructs and containing XbaI and XhoI sites (55). SRP14 was PCR amplified with p415HA_up and SRP14_low. SRP14Δ29 was PCR amplified with p415HA_up and SRP14d29_low. SRP21 was PCR amplified with p415HA_up and SRP21_low. SRP21Δ73 was PCR amplified with p415HA_up and SRP21d73_low. The PCR products were digested with XbaI and XhoI, gel purified and cloned into the equivalent sites of p413. SRP14Δ29 and SRP21Δ73 were subcloned into the NdeI and XhoI sites of pET22b. Final constructs were all verified by sequencing and are shown in Supplementary Table 2.

The sequence encoding the SCR1 RNA was amplified with an oligonucleotide containing a 5’ BamHI restriction site and the T7 promoter, and an oligonucleotide containing 3’ XhoI and DraI sites. The PCR-amplified product was cleaved with BamHI and XhoI and cloned into the equivalent sites in the pUC18 plasmid. A T7 RNA polymerase run-off transcription of the DraI-cut plasmid yielded a 522 nucleotide-long RNA with the same sequence as the endogenous SCR1 except the 5’ adenosine was replaced with a guanosine to facilitate transcription. The SCR1 deletions were similarly constructed based on the secondary model of Van Nues and Brown (34). The SCR1ΔS1 construct replaced residues 247–371 with a UUCG tetraloop, and the SCR1ΔAlu construct deleted residues 1–155 and residues 454–522. The T7 RNA polymerase run-off transcriptions of the DraI-cut plasmids yielded 401 and 303 nucleotide-long RNAs, respectively.

The actin control was a T7-promoter derivative of the previously described actin precursor RNA in the Bluescript KS(-) plasmid (93). A T7 RNA polymerase run-off transcript of EcoRI-cut plasmid yielded a 605 nucleotide-long RNA containing, from 5’ to 3’, 54 nucleotides of the plasmid, 63 nucleotides of the 5’ UTR, 10 nucleotides of the 5’ exon, 309 nucleotides of the intron, 162 nucleotides of the 3’ exon and 7 nucleotides of the plasmid.

The SRP genes (*SRP14*, *SRP21*, *SRP54*, *SRP68*, *SRP72*, *SEC65*) were PCR amplified off the pMW295 or pMW299 plasmids with oligonucleotides containing 5’ SpeI and NdeI sites and 3’ XhoI sites, and they were cloned into the SpeI and XhoI sites of the *2HA*_p424 plasmid containing an *ADH* promoter, two *HA* tags and a *CYC1* terminator (92). *SRP14* and *SRP21* also were cloned into the *GFP*_p413 plasmid. Except for *SRP68* and *SEC65*, all constructs were subcloned into the NdeI and XhoI sites of pET22b. *SRP68* and *SEC65* were re-amplified by PCR with oligonucleotides containing XhoI and BamHI sites and cloned into the equivalent sites of pET19b. *SRP101* and *SRP102* were amplified off purified chromosomal DNA using oligonucleotides with BamHI and XhoI sites and cloned into the equivalent sites of *2HA*_p424. The constructs were subcloned into the NdeI and XhoI sites of pET22b. *SRP14Δ29Cter* and *SRP21Δ73Cter* were PCR amplified with the *HA* tag and cloned into the XbaI and XhoI sites of p413 (55). *SRP14Δ29Cter* and *SRP21Δ73Cter* were subsequently subcloned into the NdeI and XhoI sites of pET22b.

The *GFP* and *MCHERRY* plasmids were constructed by amplifying genes off the pYM27-*EGFP-KanMX4* and pFA6a-*mCherry-NatNT2* plasmids, respectively. The PCR products were digested with XhoI and SalI, gel purified, and cloned into the equivalent sites of the yeast plasmids p415, p416 and p413. The *DED1*, *ded1-F162C* and *ded1-DQAD* plasmids were as previously described (13). Theses genes were subcloned into the SpeI and XhoI sites of *GFP*-p415, p414 and *MCHERRY*_p416. The *KAR2-RFP*_YIPlac204 was a gift from Benjamin Glick.

### Yeast strains and manipulations

Manipulations of yeast, including media preparation, growth conditions, transformation, and 5-FOA selection, were done according to standard procedures (94). The strains used in this study are listed in Supplementary Table 3.

The yeast GFP clone collection was purchased from Life Technologies (Ref 95702; Carlsbad, CA). The *sec61-ts* and *sec62-ts* yeast strains were a generous gift from Ron Deshaies (59). The *KAR2-RFP* strain was created by transforming the W303 (G49), *sec61* and *sec62* strains with EcoRV-linearized *KAR2-RFP*_YIPlac204 containing the N-terminus of *KAR2* (135 bp) fused to *DsRedExpress2* with the *HDEL ER*-retention sequence. The yeast *TET*-promoters Hughes Collection strains were purchased from Dharmacon (GE Healthcare, Lafayette, CO). Tetracycline-inducible strains were transformed with *GPD*-*DED1, GPD-ded1-F162C* or the empty plasmid. Cells were grown in YPD (yeast extract, peptone, dextrose) medium or in minimal medium lacking leucine (SD-LEU), serially diluted by a factor of 10 and then plated on medium with or without 10 µg/ml doxycycline.

### T7-SCR1 constructs

The oligonucleotides that were used are shown in Supplementary Table 1, where the regions of complementarity are underlined and restriction sites are shown in bold. The final constructs are listed in Supplementary Table 2.

The full-length T7-SCR1 was PCR amplified off the pMW299 plasmid (44) with the SCR1_up2 oligonucleotide containing the T7 promoter and SCR1-low oligonucleotides. The PCR product was digested with BamHI and XhoI, gel purified and cloned into the equivalent sites of pUC18. The T7-SCR1ΔAlu was constructed with the SCR1_dAlu_up oligonucleotide containing the T7 promoter and the SCR1_dAlu_low oligonucleotide. The PCR product was digested with BamHI and HindIII, gel purified and cloned into the equivalent sites of pUC18. The T7-SCR1ΔS1 construct was made as fusion PCRs with two sets of oligonucleotides. The pUC18_5’ oligonucleotide was used with SCR1_dS1_low and pUC18_3’ was used with SCR1_dS1_up. The two gel-purified PCR fragments were combined and PCR amplified with oligonucleotides pUC18_5’ and pUC18_3’. The PCR product was digested with BamHI and HindIII, gel purified and cloned into the equivalent sites of pUC18.

### Northern blot probes

The oligonucleotides used are listed in Supplementary Table 1. Oligonucleotides (150 pmoles) were 5’ ^32^P-labeled with 20 µCi of γ-^32^P ATP (3000 Ci/mmole; Hartmann Analytic) for 30 min at 37°C in 50 µl volumes with 20 units of T4 PNK (New England Biolabs) in the provided reaction buffer. Reactions were heat inactivated for 20 min at 65°C and the unincorporated radioactivity was eliminated with a G25 Illustra MicroSpin column (GE Healthcare) according to the manufacture’s instructions. Labeling efficiency was determined by comparing the radioactivity of the recovered material with that retained on the column. Blots were incubated with 15 pmoles of probes overnight at 42°C. Blots were washed two times at 42°C with 2X SSC buffer (Euromedex, Souffelweyersheim, France) with 0.1% SDS added, washed two times with 0.2X SSC buffer with 0.1% SDS, dried and then subjected to autoradiography with a BAS-MS imaging plate (Fujifilm) overnight. Exposures were revealed with a Typhoon FLA9500 phosphoimager (GE Healthcare).

### *In situ* localization

To analyze the location of Ded1 relative to the ER or SRP proteins, we first used the green fluorescent protein (GFP)-tagged *ded1-DQAD* plasmid that was transformed into *sec61-ts* and *sec62-ts* mutant strains with the integrated *KAR2-RFP* plasmid (Supplementary Tables 2 and 3). Cells were grown in SD-LEU to an OD_600_ of ∼0.9–1.0 (logarithmic phase) at 24°C and then shifted to 37°C for 15 min. We subsequently used mCherry-tagged *ded1-DQAD* plasmid that was transformed into GFP-tagged *SRP14* and *SRP21* expressed from the chromosome (Supplementary Tables 2 and 3). Cells were grown in SD-LEU to an OD_600_ of 0.95 at 30°C. Finally, GFP-tagged *SRP14* or *SRP21* and mCherry-tagged *ded1-DQAD* plasmids were transformed in the *xpo1-T539C* strain (70). Cells were grown in minimal medium lacking histidine and uracil (SD-HIS-URA) to an OD_600_ of 0.4 (early logarithmic phase) at 30°C, and then they were split into two parts: one-half was resuspended in SD-HIS-URA and the other in SD-HIS-URA supplemented with ∼200 nM leptomycin for 1 h.

Fluorescence microscopy was done with a Zeiss Observer.Z1 microscope (Carl Zeiss Microscopy GmbH) with a 63x oil immersion objective equipped with the following filter sets: Alexa 489, filter set 10 from Zeiss for GFP (Excitation BP 450-490, Beam Splitter FT 510, Emission BP 515/65), HC-mCherry, filter set F36-508 from AHF for mCherry and RFP (Excitation BP 562/40, Beam Splitter 593, Emission BP 641/75). Images were acquired with a SCMOS ORCA FLASH 4.0 charge-coupled device camera (Hamamatsu photonics) using the Zen 2012 Package Acquisition/Analysis software and processed with Zen 2012 and Photoshop CS3.

### RNA transcripts

RNAs were produced as run-off transcripts using T7 RNA polymerase and the MEGAscript kit (Ambion Thermo Fisher Scientific, Waltham, MA) according to the manufacture’s instructions. In brief, reactions were done in 20–40 µl volumes with 1–2 µg of linearized DNA and incubated for 5–6 h at 37°C. The template was then digested with TURBO DNase, multiple reactions were combined, the solution diluted to 500 µl final with high salt buffer (300 mM potassium acetate, 50 mM Tris-base, pH 8.0, 0.1 mM EDTA), extracted with an equal volume of water-saturated phenol (MP Biomedicals, CA), extracted with an equal volume of chloroform-isoamyl alcohol (24:1) and precipitated overnight at −20°C with 2.5–3 volumes of ethanol. The RNA was recovered by centrifugation in an Eppendorf 5415R at high speed at 4°C for 15 min. The supernatant was discarded, the pellet washed with 300 µl of cold 70% ethanol, and the pellet dried in a SpinVac (Savant Thermo Fisher Scientific) for 20 min. To eliminate trace contaminates, the RNA was resuspended in 400 µl high salt buffer and re-ethanol precipitated at −20°C. The pellet was recovered, washed and dried. The final pellet was resuspended in 20 mM Tris-base, pH 7.5, 0.1 mM EDTA and stored at −20°C until needed. The concentration was determined using an absorbance at 260 nm of 32 µg/ml/cm, which was based on the calculated values of Oligo 7 software (Molecular Biology Insights, Inc, Colorado Springs, CO).

### Recombinant protein expression and purification

Recombinant Ded1-His was expressed from the pET22b plasmid (Novagen) and purified as previously described (46). SRP His-tagged proteins were transformed into the Rosetta (DE3) *E. coli* strain. Cultures containing 500 ml of cells at OD_600_ of 0.5 were induced with 0.5 mM isopropyl-1-thio-ß-D-galactopyranoside (IPTG) for 1 h at 37°C for SRP14, Sec65 and SRP101; 16 h at 16°C for SRP68 and SRP72; and for 2 h at 30°C for SRP54 and SRP21. Cells were collected by centrifugation and the pellets were resuspended in 5 ml of lysis buffer containing 20 mM Tris-base, pH 8.0, 0.5 M NaCl and the following protein-specific conditions: 20 mM imidazole, 8 mM ß-mercaptoethanol and 1% Triton X-100 for SRP14, SRP68, SP101 and Sec65; 20 mM imidazole and 3 mM ß-mercaptoethanol for SRP21; 10 mM imidazole and 3 mM ß-mercaptoethanol for SRP54; and 20 mM imidazole and 1mM ß-mercaptoethanol for SRP72. The equivalent of 1 mg/ml lysozyme was added for each condition, the cells kept on ice for 30 min and then the cells were lysed by sonication at 4°C until the lysate became clear. The material was centrifuged for 40 min at 14,000 rpm in a JA-12 rotor (Beckman Coulter, Brea, CA) at 4°C and the supernatant was loaded onto a 1 ml nickel nitrilotriacetic acid-agarose column (Ni-NTA, Qiagen, Hilden, Germany) previously equilibrated with the corresponding lysis buffer. After two washes, the SRP His-tagged proteins were eluted with lysis buffer containing 150 mM imidazole. Purified proteins were supplemented in 50% glycerol and were quantified using the Bradford protein assay kit (Bio-Rad). Proteins were aliquoted and stored at −80°C until needed. Recombinant purified proteins, supplemented with SDS sample buffer, were resolved by 12% SDS polyacrylamide gel (SDS-PAGE) and stained with Coomasie Blue (Instant Blue). The properties of the different proteins are shown summarized in Table 3 and the purified proteins shown in Supplementary Figure 1.

### Immunoglobulin G-protein A Sepharose-bead pull-down experiments

The G50 yeast strain was transformed individually with *2HA*_p424 plasmids containing two amino-terminal HA tags and the genes for *SRP14, SRP21, SRP54, SEC65, SRP68, SRP72* and *SRP101*. Cells were grown to an OD_600_ of 0.8–1 in minimal medium lacking tryptophan (SD-TRP), recovered by centrifugation, washed with cold water, frozen in liquid nitrogen and stored at −80°C until needed. The cells were resuspended in an equal volume of lysis buffer containing 20 mM HEPES, pH 7.4, 150 mM NaCl, 5 mM MgCl_2_, 0.1 mM EDTA, 5 mM DTT and 1X protease inhibitor cocktail (Roche cOmplete EDTA-Free). An equal volume of 425– 600 µm glass beads (Sigma-Aldrich, St. Louis, MI) was added and the cells were broken by vortexing in a FastPrep-24 (M.P. Biomedicals) at 4°C with four cycles of 30 sec and 5 min rests. The cell debris was removed by centrifugation for 5 min at 6,000 rpm in an Eppendorf 5415R centrifuge at 4°C. The lysates were further clarified by centrifugation two to three times at 13,000 rpm for 10 min each at 4°C. The protein concentrations were determined with a Bio-Rad Protein Assay kit according to the manufacture’s instructions using bovine serum albumin (BSA) as a standard.

Protein A-Sepharose CL-4B beads (GE Healthcare) were prepared by first washing them twice in IPP150 buffer containing 20 mM Tris-base, pH 7.4, 150 mM NaCl, 0.1% Triton-X100, and 1 mM MgCl_2_. Then, 50 µl of beads was incubated overnight with mixing in 10 volumes of IPP150 buffer at 4°C with 0.4 mg/ml BSA, 0.4 mg/ml heparin, and 20 µl of serum containing Ded1, SRP21, HA (Covalab, Bron, France) or pre-immune immunoglobin G (IgG). The beads were washed three times with 800 µl of 1X PBS, and then mixed at 4°C for 2 h with G50 extracts containing 300 µg of protein and 10 volumes of 1X PBS buffer supplemented with BSA and heparin. The beads were washed three times with 800 µl of 1X PBS, and the bound proteins eluted twice with 300 µl of 0.1 M glycine, pH 2.3 for 15 min at 4°C with mixing. The pH of the eluted proteins was then adjusted to pH ∼7 with NaOH.

### Other pull down experiments

Protein A-Sepharose beads were prepared as described above. Ded1 or SRP21 IgG were crosslinked to beads with 0.2% glutaraldehyde as described previously (95) and were rigorously washed with 1X PBS. The equivalent of 4–6 µg of purified Ded1 and SRP proteins were incubated with 300 µl 1X PSB supplemented with 2 mM MgCl_2_ for 45 min at 30°C. Fifteen µl of Protein A-Sepharose beads were directly added to the protein mixture and mixed by rotation for 1 h at 18°C. Prior to elution, beads were washed three times with 1X PBS. The bound proteins were eluted with 30 µl glycine, pH 2.3, for 15 min at 4°C on a rotating wheel platform. The acidity of the reaction was neutralized by adding NaOH. Co-immunoprecipitated purified proteins, supplemented with SDS sample buffer, were resolved on a 12% SDS-PAGE and stained with Coomasie Blue.

### Western blot analysis

The eluted proteins were concentrated by making the solution 150 µg/ml in sodium deoxycholate and the proteins precipitated with 15%, final, of trichloroacetic acid (TCA) for 16 h at 4°C. The solution was centrifuged 30 min in an Eppendorf 5415R centrifuge at high speed, and the recovered pellet washed with cold acetone, dried and resuspended in loading buffer containing 50 mM Tris-base, pH 6.8, 2% sodium dodecyl sulfate (SDS), 1% β-mercaptoethanol, 0.02% bromophenol blue and 10% glycerol. The eluted proteins were separated on SDS-Laemmli gels, transferred to Amersham Protran nitrocellulose membranes (GE Healthcare Life Science) and probed with primary IgG against Ded1, SRP21, Sec65 or HA. The anti-Sec65 antibody was a generous gift from Martin R. Pool. Horseradish peroxidase-conjugated anti-rabbit (for Ded1, Sec65, SRP21; Covalab) and anti-mouse (for HA; Covalab) were used as secondary antibodies, and the signals were revealed with a Clarity Western ECL Substrate kit (Bio-Rad, Hercules, CA) using a Bio-Rad ChemiDoc XRS+ and Image Lab 5.2 software. The Ded1-IgG and SRP21-IgG were produced by Covalab using purified recombinant Ded1 or SRP21, respectively.

### Polysome sucrose gradients

G50 yeast strains containing HA-tagged SRP proteins were grown to an OD_600_ of 0.8–1.0 in SD-TRP, cycloheximide (Sigma Aldrich) was added to a final concentration of 50 µg/ml, and the cells were incubated on ice for 10 min. Cells were harvested by centrifugation, washed with a lysis buffer containing 10 mM Tris-base, pH 7.4, 100 mM NaCl, 5 mM MgCl_2_, 0.1 mg/ml cycloheximide, 1 mM DTT and 1X cOmplete EDTA-free protease inhibitor cocktail (Roche), the pelleted cells were then frozen in liquid nitrogen and stored at −80°C until needed.

Cells were resuspended in an equal volume of lysis buffer, one volume of 425–600 µm glass beads was added, and then the cells were lysed in a FastPrep-24 as described above. Cell debris was removed by centrifugation in a Beckman JA-12 rotor at 6000 rpm for 5 min at 4°C, and then the lysates were further clarified by centrifugation in an Eppendorf 5415R centrifuge at 13,000 rpm for 10 min at 4°C. The equivalent of 200 µl of cell extract with an OD_260_ of 12 was loaded onto a 10–50% sucrose gradient and centrifuged in a SW41 rotor (Beckman Coulter) at 39,000 rpm for 2.45 h at 4°C. Half milliliter fractions from the gradient were collected with a Retriever 500 (ISCO) fraction collector and monitored with a UV-6 UV/VIS detector (ISCO) at 254 nm. Fractions were subsequently split in half for either RNA extraction or Western blot analysis.

The protein fractions were concentrated by adding 150 µg/ml, final, of sodium deoxycholate and then the solutions were made 15% in TCA, centrifuged in an Eppendorf 5415R, the recovered pellets were washed twice with cold acetone, and then the protein pellets were dried. The recovered material was resuspended in SDS loading buffer, separated by electrophoresis on a 12% SDS-PAGE gel and electrophoretically transferred to nitrocellulose membranes. The Western blots were undertaken as described above.

The RNA fractions were made 0.3 M in potassium acetate, extracted with an equal volume of water-saturated phenol, extracted twice with an equal volume of chloroform-isoamyl alcohol (24:1), and then ethanol precipitated overnight at −20°C. The RNA was recovered by centrifugation, dried, resuspended in 1X RNA loading buffer (Thermo Scientific), and then electrophoretically separated on a 6% polyacrylamide gel containing 7 M urea and ethidium bromide (to reveal 18S and 23S rRNAs). The RNA were subsequently electrophoretically transferred to Amersham Hybond-N+ nylon membranes (GE Healthcare) and probed with a ^32^P-labeled DNA oligonucleotide specific for SCR1 (Supplementary Table 1). The image was visualized with a Typhoon FLA9500 phosphoimager (GE Healthcare).

### Reverse-transcriptase PCR

Samples were digested with Proteinase K by adding 1 mg/ml proteinase K (Sigma #P2308-100MG), 1% triton X-100, 0.5% SDS, 5 mM CaCl_2_ in an Eppendorf Thermomixer Comfort (15 sec 1000 rpm, 90 sec rest) at 55°C for 35 min. Total RNA (input condition) and RNA from the eluate were recovered by the addition of 0.3 M potassium acetate, extracted with an equal volume of water-saturated phenol, extracted twice with an equal volume of chloroform-isoamyl alcohol (24:1), and then ethanol precipitated overnight at −20°C. The RNA pellets from the ethanol precipitations were resuspended in 20 µl nuclease-free water. RNA was reverse transcribed with the Superscript III kit (Invitrogen, Carlsbad, CA) according to the manufacture’s instructions. In brief, 4.6 µl of the resuspended Ded1, pre-immune IgG pull-downs, and 0.5 µg of total yeast RNA were combined with 1 pmole of the 3’ primers specific for SCR1, PGK1 or RPL20B RNAs (Supplementary Table 1). The reactions were heated to 50°C for 5 min and then 10 mM DTT, 0.75 µl of Superscript III Reverse Transcriptase (RT), 1X final of RT buffer and 3 mM final of dNTPs were added. The reactions were incubated for 90 min at 50°C. The RNAs were hydrolyzed by adding 40 µl of a solution with 150 mM KOH, 20 mM Tris-base and incubating at 90°C for 10 min. The solution was neutralized by adding 40 µl of 150 mM HCl. The PCR amplification was performed with 10 µl of the reverse transcriptase product in a 50 µl PCR mix containing 1 unit of Phusion High-Fidelity DNA polymerase (New England Biolabs, Évry-Courcouronnes, France), 1X HF Phusion buffer, 0.2 mM dNTPs, and 0.5 pmoles of the respective gene-specific 5’ and 3’ primers (Supplementary Table 1). PCR reactions were done for 25 cycles in a Bio-Rad T100 Thermal Cycler. PCR products were purified with a NucleoSpin Gel and PCR Clean-up kit (Macherey-Nagel, Düren, Germany), eluted with 50 µl elution buffer and 5 µl aliquots were analyzed with agarose loading buffer on a 2% agarose gel containing ethidium bromide

### *In vitro* RNA-dependent ATPase activities

The ATPase assays were based on a colorimetric assay using molybdate-Malachite green as previously described (46). We typically used 23 nM of SCR1 or actin RNA and 0.14 µg/µl of whole yeast RNA (Roche) that was purified on a DEAE-Sepharose column to remove inhibitors. For the latter RNA, fractions from an elution with increasing concentrations of NaCl were assayed with purified Ded1, and the most active fractions were combined, concentrated by ethanol precipitation and subsequently used in the assays. The poly(A) RNA was from Sigma. Assays were performed in a reaction mix with 20 mM Tris-base, pH 7.5, 50 mM potassium acetate, 5 mM magnesium acetate, 0.1 µg/µl BSA, and 2 mM DTT. Purified proteins and RNAs were pre-incubated for 30 min at 30°C to equilibrate the different components. We used the Ded1-K192A (GAT) mutant in motif I as a negative control as it had no detectable ATPase activity (46). Reactions were started by adding 1 mM final of ATP and taking aliquots over the time course. The reactions were stopped by making the solutions approximately 60 mM final in EDTA. The Malachite green solution was added as previously described (46) and the absorption measured with a Tecan NanoQuant Infinite M200Pro microtiter plate reader at 630 nm. Enzymatic reaction velocities were determined by a linear regression fit over the initial linear phase of the reaction with five data points over a time course of 45 min using optimized protein concentrations. We used a serial dilution of Phosphate Phosphorous Standard for IC (Fluka Sigma-Aldrich) for each experiment to determine the corresponding phosphate concentration from the absorption. Experimental data were analyzed with Kaleidagraph 4.5.2 software (Synergy, Reading PA).

### Electrophoretic mobility-shift assays

An EMSA-agarose technique was used as previously described with minor modifications (96). Briefly, the assay were performed with 0.150 µM SCR1 or actin RNA and variable concentrations of the indicated proteins, which were incubated together in 1X EMSA buffer (20 mM Tris-base, pH 8.8, 70 mM KOAc, 2 mM MgCl_2_, 10 µg/µl BSA, 1 mM DTT) for 15 min at 30°C in the presence or absence of 5 mM AMP-PNP or ATP in a volume of 8 µl. Two µl of 30% glycerol was added to the samples, and they were loaded onto 0.75 mm thick, 1% agarose (Molecular Biology Grade) gels containing 1X TBE buffer (45 mM Tris-base, 45 mM boric acid, pH 8.8, and 2 mM EDTA; Sigma) and ∼0.016 µg/ml ethidium bromide. Gels were run in a mini-plus horizontal electrophoresis (Scie-Plas) in 1X TBE buffer at 220V for ∼13 min at 4°C and were imaged with a Gel Doc XR+ (Bio-RAD) and Quantity One 4.6.9 software (Bio-Rad).

## ACKNOWLEDGMENTS

We thank Caroline Lacoux and Stéphanie Ørum for helpful discussions, Martin R Pool for the Sec65 IgG and for the pMW295 and pMW299 SRP plasmids, Benjamin Glick for the *KAR2-RFP*_YIPlac204, Jiří Hašek for the pFA6a-mCherry-NatNT2 plasmid, Michael Lisby and Rodney Rothstein for the G49 and G50 yeast strains, and Claude Thermes, Yves Daubenton, Erwin Van Dijk, Yan Jaszczyszyn, Pauline Francois, Marina Cavaiuolo and Benoist Laurent for help with RNAseq.

## Supplementary Files

**Supplementary Table 1.**
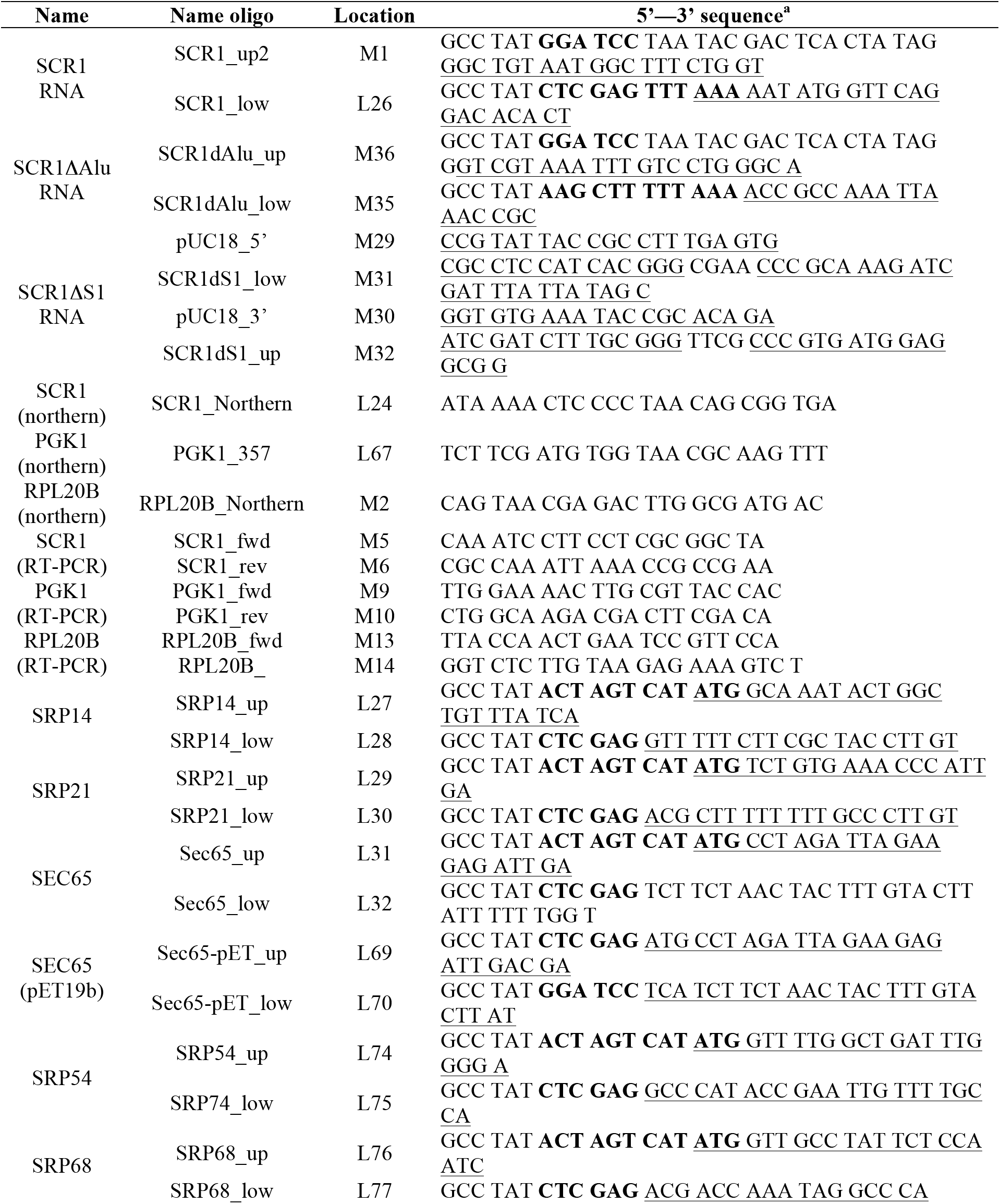
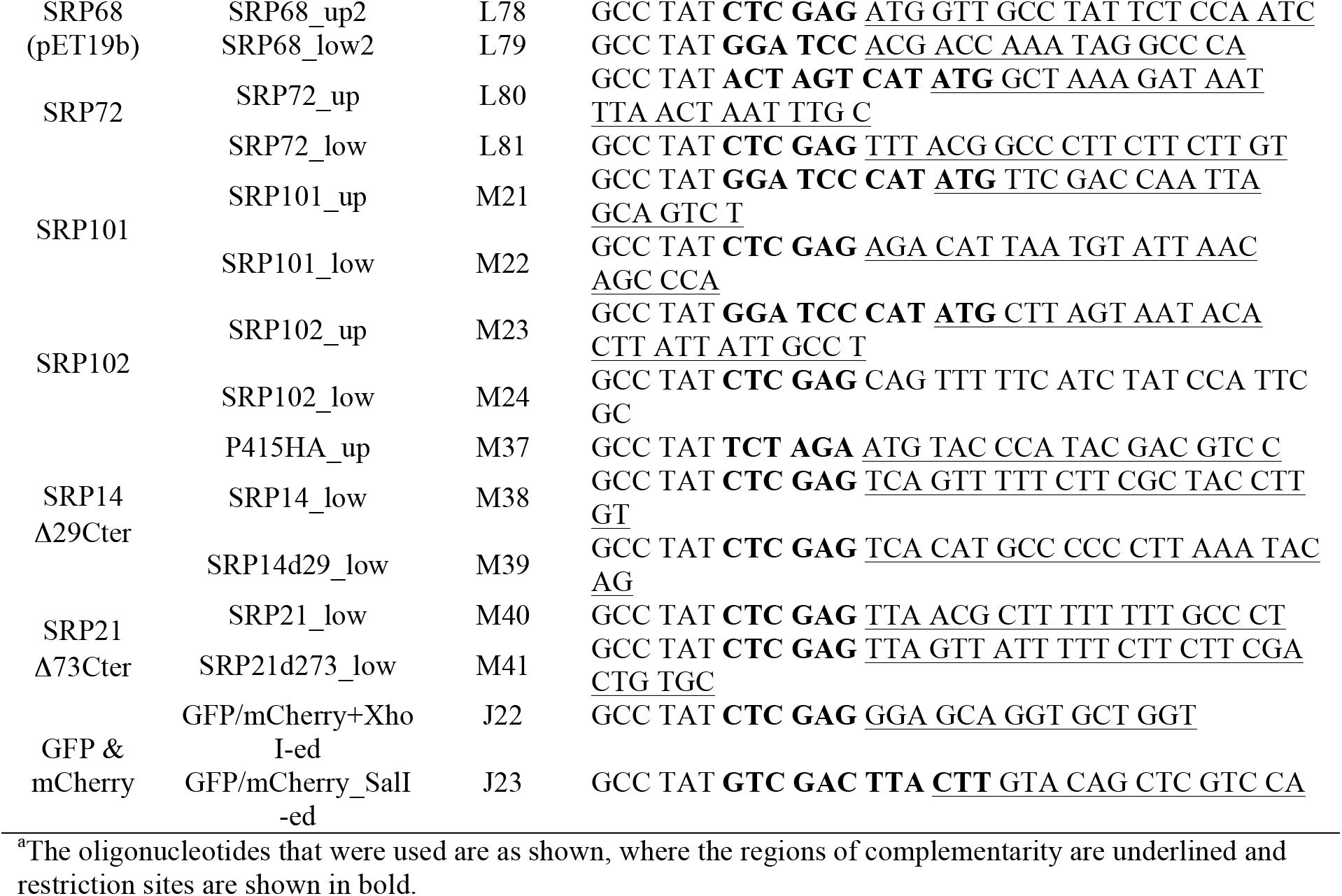
Oligonucleotides used in this study

**Supplementary Table 2.**
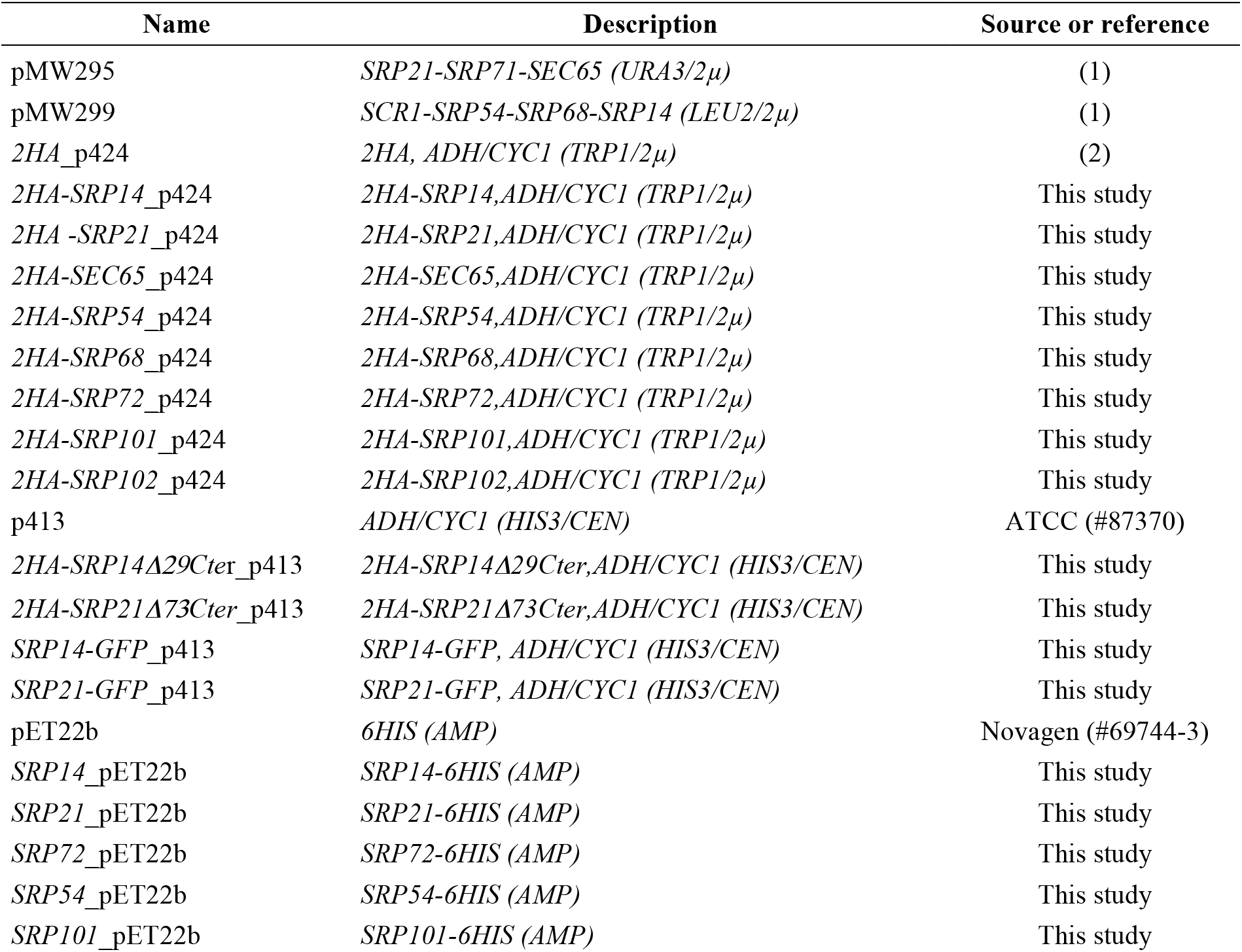
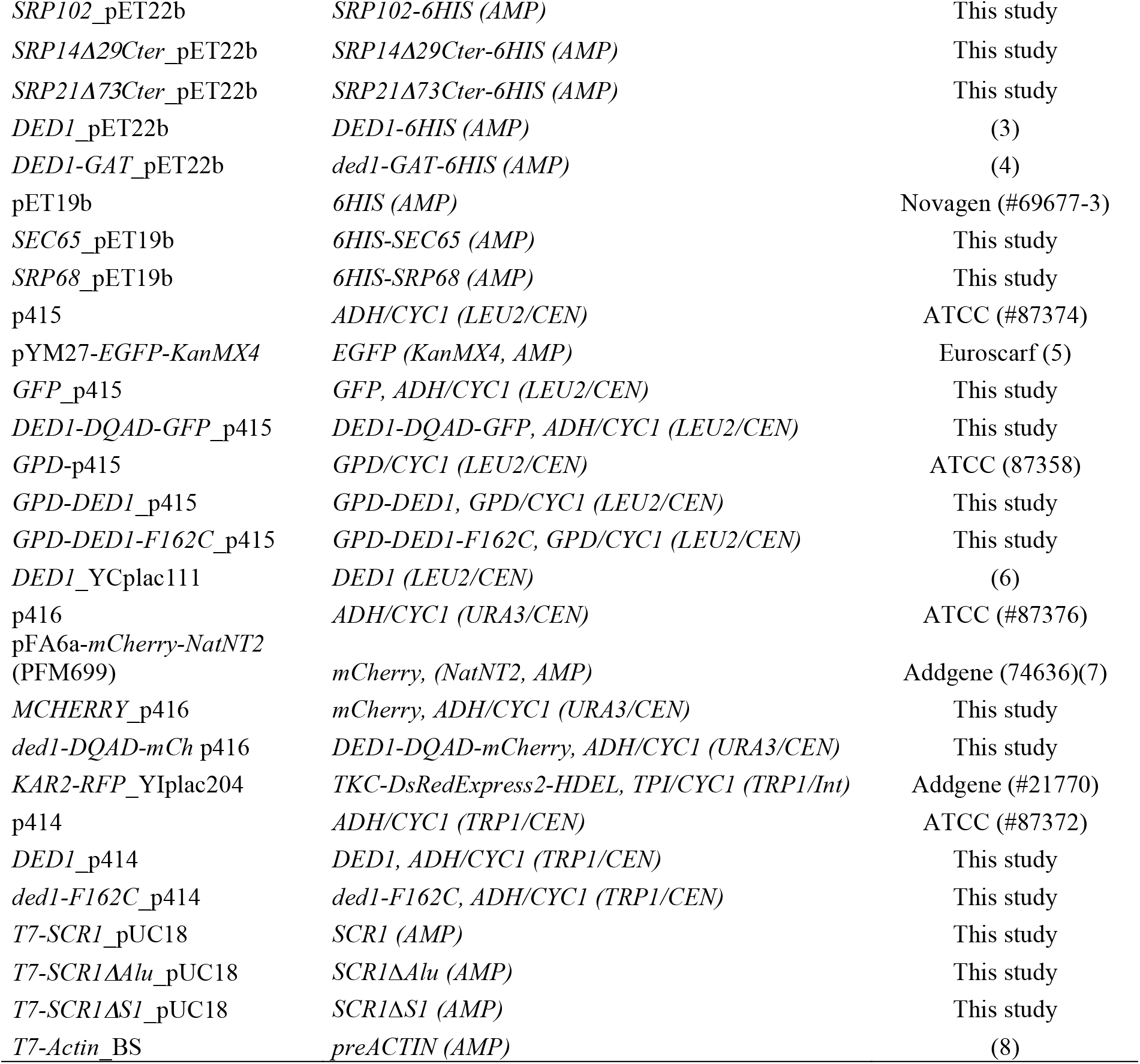
Constructs used in this study

**Supplementary Table 3.**
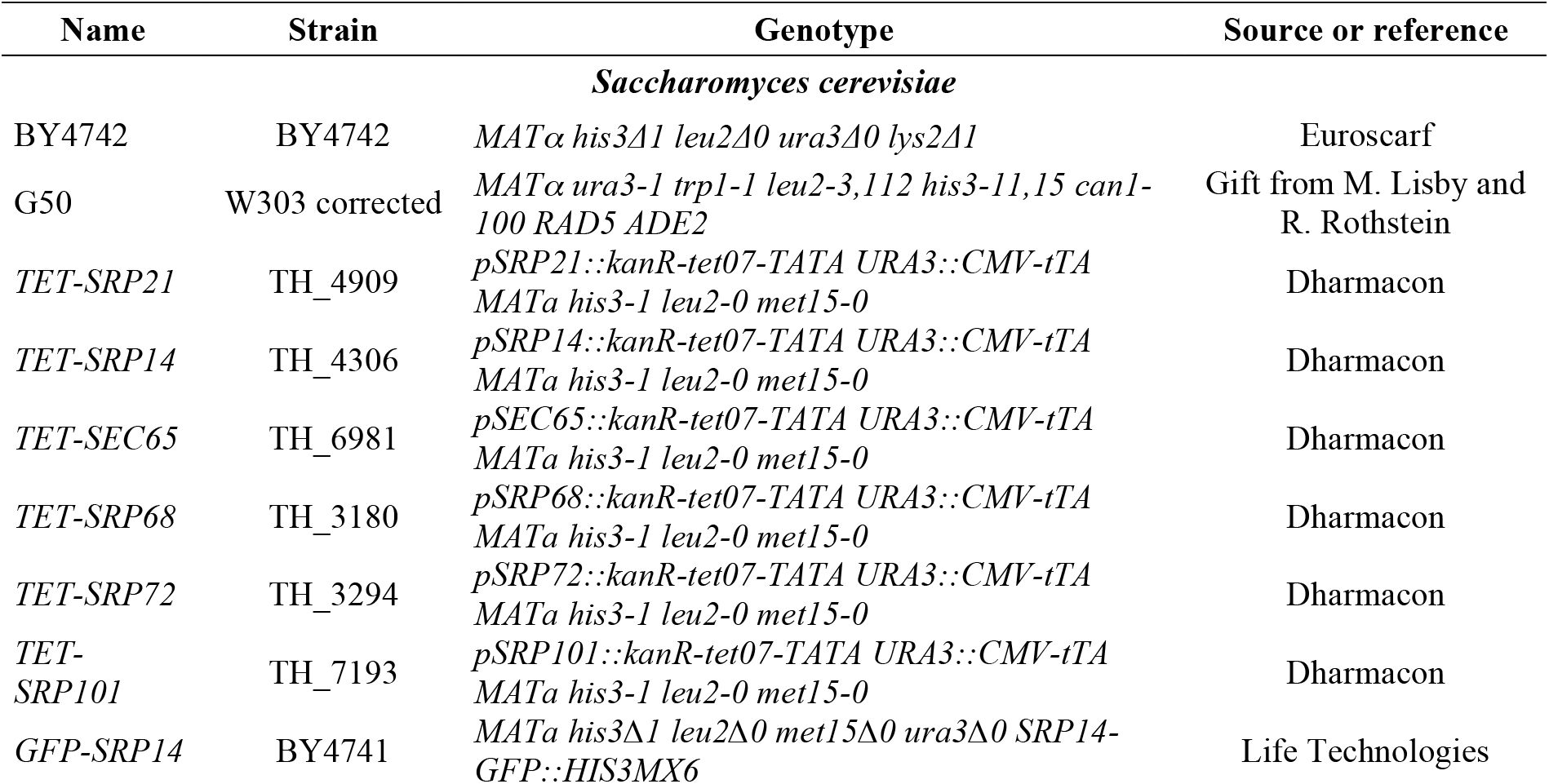
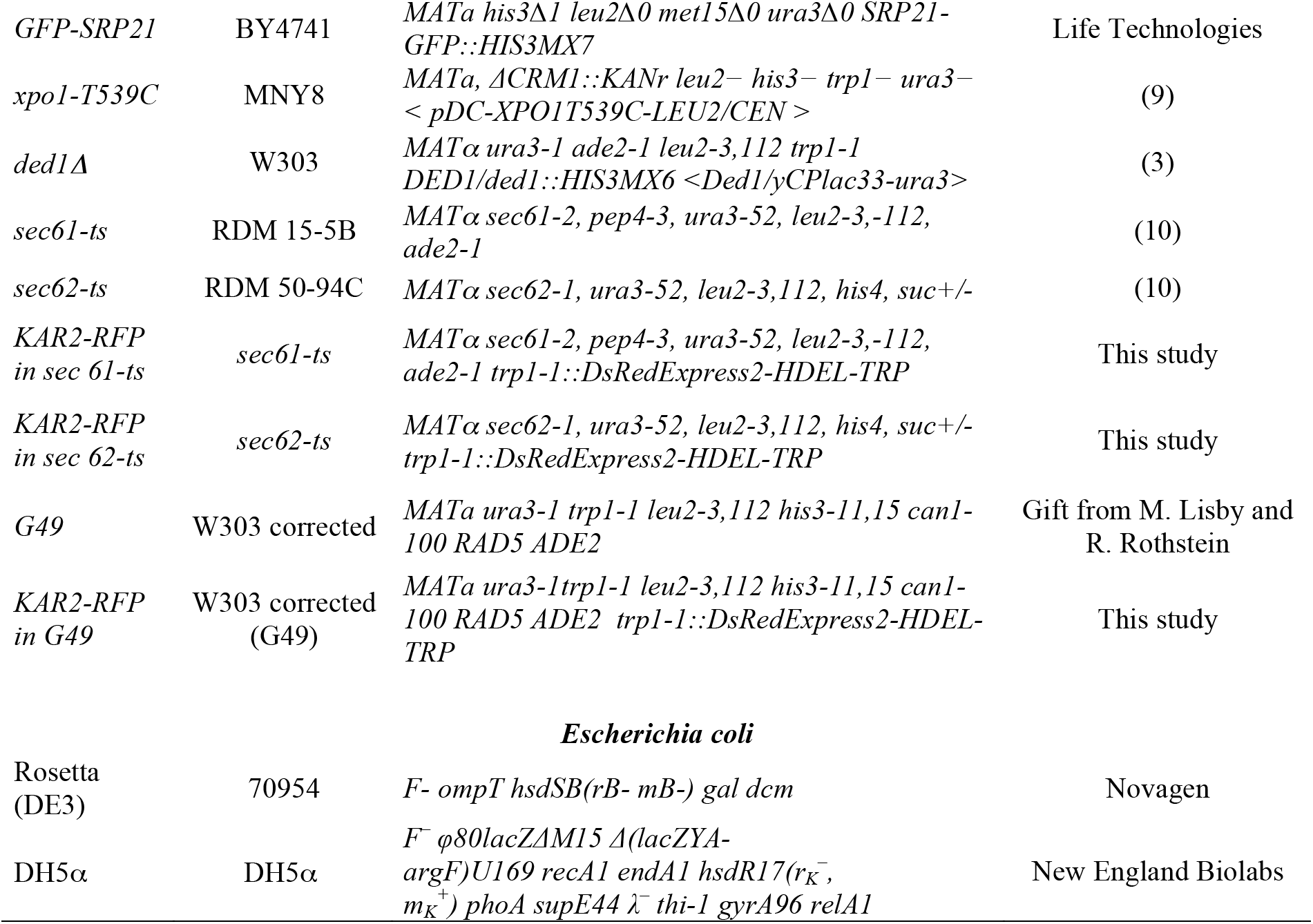
Yeast and bacterial strains used in this study

**Supplementary Figure 1.**
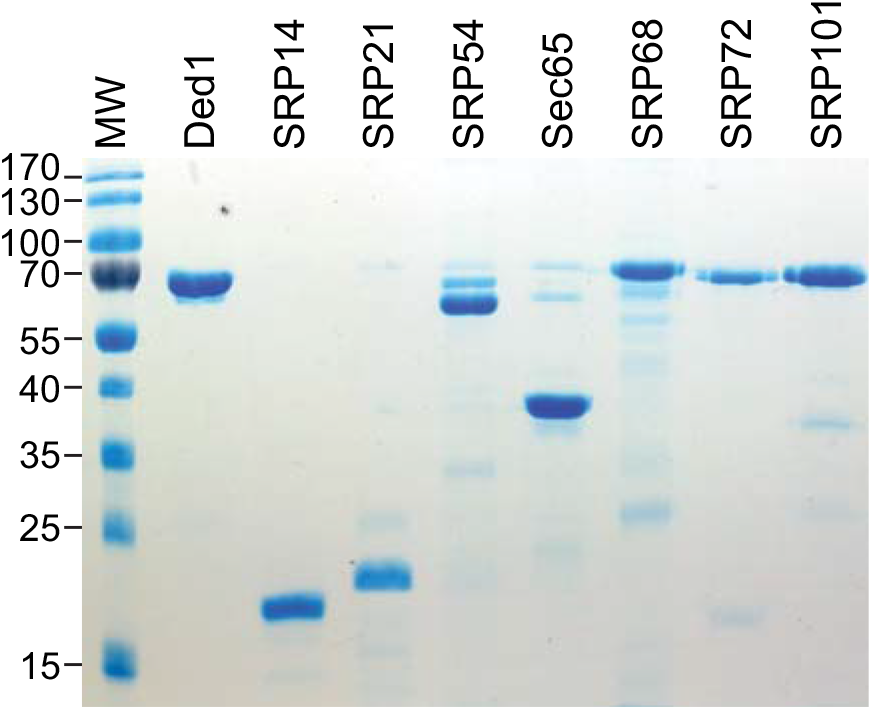
Purified His-tagged recombinant proteins. Aliquots of 1.3 µg of proteins purified on a Ni-NTA column were electrophoretically separated on a 12% PAGE and stained with Coomassie blue.

